# Spatiotemporal transcriptomic niches of complement pathway and serine protease inhibitor activation in aging and infection

**DOI:** 10.1101/2024.11.04.621811

**Authors:** Fabian Kern, Viktoria Wagner, Vanessa Wahl, Nicole Ludwig, Simon Graf, Jeremy Amand, Aulden Foltz, Micaiah Atkins, Blen Kedir, Alexandros Hadjilaou, Philipp Donate, Matthias Flotho, Friederike Grandke, Shusruto Rishik, Philipp Wartenberg, Monika I. Hollenhorst, Daniel Krug, Hanna Lotter, Rolf Müller, Ulrich Boehm, Oliver Hahn, Thomas Jacobs, Gabriela Krasteva-Christ, Tony Wyss-Coray, Andreas Keller

## Abstract

Aging is a multifactorial and complex physiological process, affecting every organ with characteristic manifestations. Understanding the molecular mechanisms that drive aging processes is crucial to targeting age-related disorders. Recent reports suggest that severe post-infection syndromes can partially accelerate aging. However, the underlying gene-encoded regulatory interplay, whether being shared or distinct between aging and infection biology are poorly understood. Here, we employed spatial transcriptomics to establish a multi-organ atlas (brain, heart, kidney, liver, lung, and spleen) across the mouse lifespan (4, 17, and 26 months). Dissecting high-quality fresh-frozen tissue samples at unbiased molecular resolution, we found both organ-specific and cross-organ gene dysregulation upon aging. We identified age-related trajectories in gene expression and cell state, some only detectable within their spatial context, and provide validation at subcellular resolution. The most prominent effect was organ-wide immune system activation with spatially variable severity. We therefore evaluated how aging mimics the expression signatures observed in systemic infection, using spatial transcriptomics slices from young mice infected with Plasmodium berghei ANKA. While on the gene level the effect sizes caused by the infection outweighed those of aging, we reveal a shared activation of the early complement pathway (C4b) and serine protease inhibitors (Serpin gene family) within by phenotype distinct spatial niches. We show that this common RNA signature is driven by tissue-specific cell types and eventually affects protein levels in the aged brain, rendering them a target for future mechanistic and drug discovery studies. Taken together, our study provides a coherent in-depth and cross-organ transcriptomics atlas to systematically study aging and infection in the mouse at spatiotemporal resolution.

**Key highlights:**

- Large-scale and high-resolution atlas of spatial transcriptomics from six organs to study aging and systemic infection across two mouse cohorts.
- Strong transcriptional alterations found in distinct organ-specific niches for aging and acute malaria, with organ- and cell type-associated immune responses.
- Dysregulation of early complement proteases (C4b) and serine protease inhibitors (Serpina3n) as common theme across central nervous system and peripheral organs.

## Main

The population-wide risk of cardiovascular, neurodegenerative, and metabolic syndromes is inextricably connected to societal aging, which involves an organism-wide deterioration of tissue structure and function^1^. Previous genomics studies across multiple organs have examined molecular changes associated with aging and provided insights into unique trajectories occurring in both organs and individual cell types^2–5^. Single-cell RNA-seq approaches revolutionized our understanding of cell type-specific gene regulation in health and disease. The human and the murine brain have been of particular interest for the application of -omics approaches to identify molecular mechanisms in age-related cognitive decline and progressive neurodegeneration^6–9^. However, a detailed and spatially-resolved understanding of the molecular programs underlying the functional consequences of aging in the hundreds of mammalian cell types within and between organs is still missing. Recent advances in image- and sequencing-based spatial transcriptomics approaches have enabled us to profile the expression of thousands of genes from tissue slices in two- or three-dimensional space^10^. These approaches facilitate the discovery of fine-grained structural changes across tissue sections. Thereby, they overcome the existing limitations of single-cell RNA-seq approaches that typically miss the microenvironment, imposed by the inevitable tissue dissociation process leading to the loss of spatial information^11^. Among those, unbiased genome-wide approaches operating at the (super-)cellular level together with targeted, gene panel-based approaches of sub-cellular resolution have been developed. In the context of aging, spatial transcriptomics hold the promise of narrowing down the cell etiology, specifically by identifying early cellular hot spots of gene dysregulation^9^.

The mechanistic interplay between the innate and adaptive immune systems in aging, for instance the observed chronic low-grade sterile inflammation (“inflammaging”), has been a long withstanding aspect in the field^12^. But, the task to discern cause (driver) from effect (passenger) of the aging process is still one of the major ongoing challenges, in particular the aspect how acute events might eventually promote chronic symptoms. In fact, an entire spectrum of inflammatory phenotypes of increasing severity exists, although their relationship to human aging is mostly unclear. The wealth of COVID-19 accelerated interdisciplinary research has pinpointed a potential causal connection between severe infectious diseases and aging-associated disease development on a global scale^13^. This innumerable data indicates common molecular immune pathways across the human body underlying both systemic aging and infection. Still, when and where such a potentially conserved biological mechanism is at play has remained unclear. Therefore, we performed 168 spatial transcriptomics experiments at an unprecedented scale utilizing two independent mouse cohorts, reporting 100 manually annotated spatial compartments based on genome-wide expression clustering of 383,040 high-quality barcoded sequencing spots. For in-depth validation of brain aging signatures, we used STOmics Stereo-seq, an unbiased genome-wide spatial transcriptomics technique, providing sub-cellular views on gene expression. We comprehensively compared organismal aging against systemic infection on the RNA level, both in the periphery and the central nervous system, and identified a shared mechanism of complement dysregulation occurring in specific local niches.

## Spatial transcriptomics of aging mouse organs

We first performed spatial transcriptomics experiments using the spot-based 10x Visium platform on a wildtype mouse aging cohort, consisting of up to five mice per timepoint (young: 4 months; middle-aged: 17 months; aged: 26 months) followed by deep sequencing (**Fig. 1a**). For each individual mouse, we processed single sections for the peripheral organs (heart, kidney, liver, lung and spleen) and five sections from the brain (Bregma: –1.7, -2.06, -2.7, -3.52 and -4.72). Following a stringent image-based and sequencing metrics-based quality control, we focused our downstream analysis on 111 high-quality samples (**Extended Data Fig. 1-2**, **Supplementary Table 1**). With a tissue-specific average number, out of a total of 4992 spots available per capture area, we maximized the number of spots located under tissue (mean of 2522 before QC, 2441 after QC), while retaining a high level of reproducibility between replicates of the same organ and across all ages (**Fig. 1b**). The data set covered around 22.3 billion reads (∼201 Mio. per sample) for an average of 3783 median genes detected at each spot (12,048 average of median UMIs). To this end, we selected histologically motivated cutting angles, trying to optimize capture of the inherent cellular heterogeneity of each peripheral organ (**Fig. 1c**). After filtering individual spot locations of lower quality (cf. Methods), we found specific properties of the spot clustering in our cleaned data set. While gene expression programs in spots of the structurally more defined organs like the kidney primarily clustered according to their spatial domain, the spot clustering in more homogenous tissues was overly driven by cell type-specific markers (**Fig. 1d, Supplementary Table 2**). In fact, the size and number of distinct expression clusters correlated with histological tissue complexity. We thus followed a hierarchical annotation approach, defining either a local region of gene expression (e.g. “Atria” in the heart), or if possible, naming each cluster after the most enriched cell type (e.g. “Periportal hepatocytes” in the liver) based on known marker genes.

**Figure 1:**
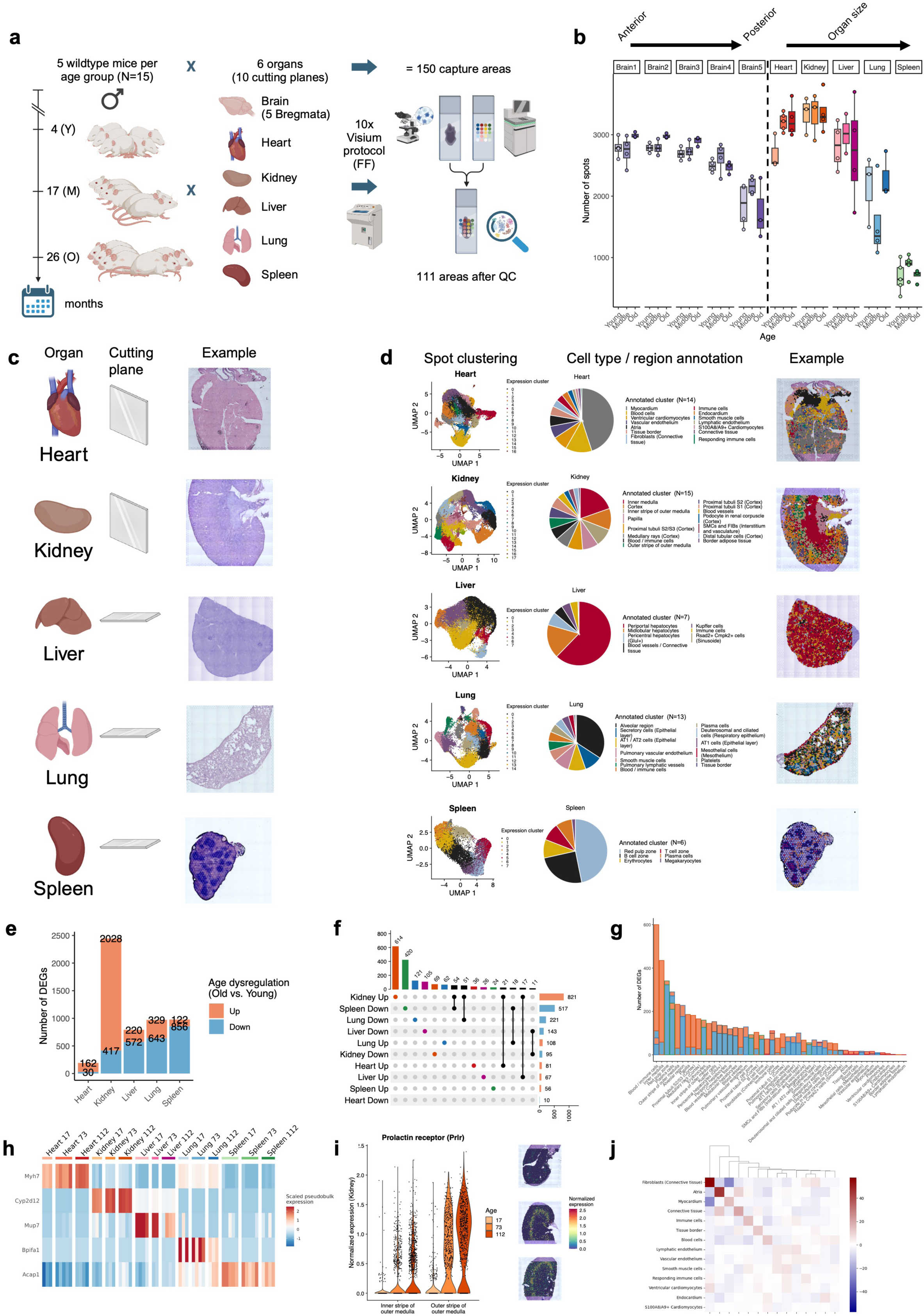
Spatial transcriptomics of the aging mouse. **a**, Illustration of the experimental workflow conducted for the mouse aging cohort. **b**, Distribution of the number of cleaned spots under tissue per organ and age. Average number of cleaned spots located under tissue varied between organs (Brain1: 2825, Brain2: 2839, Brain3: 2764, Brain4: 2534, Brain5: 1939, Heart: 3093, Kidney: 3342, Liver: 2824, Lung: 1926, Spleen: 759), out of the 4992 spots possible per capture area. In general, the number of spots per sample was influenced by slight inter-individual variances in organ size. Colored boxes span the first to the third quartile with the line inside the box representing the median value. The whiskers show the minimum and maximum values or values up to 1.5-times the interquartile range below or above the first or third quartile if outliers are present (shown as separate, colored dots). **c**, Illustration of cut angle plane and one representative H&E-stained capture area per peripheral organ. **d**, From left to right: Two-dimensional UMAP representation of colored spot clusters computationally integrated by organ (top to bottom), pie chart showing the proportion of annotated clusters across all samples, one representative annotated Visium sample with the spot cluster identities plotted over the H&E-stained tissue image. **e**, Number of differentially expressed genes per aging peripheral organ (old vs. young) and direction of dysregulation. Total DEG counts were derived across all organ clusters, without removing duplicates. **f**, UpSet plot comparing the unique gene sets across all organ cluster DEGs shown in (**e**). **g**, Bar plot showing the number of DEGs per annotated spot cluster. Tissues sharing the same cluster names are stacked on top of each other. **h**, Heatmap showing scaled expression of representative aging DEGs (rows) across all pseudobulk samples (columns). **i**, Normalized expression of Prolactin receptor (Prlr) in the kidney across two spot clusters and three aging groups (left) and representative spatial transcriptomics samples from each aging group with Prlr expression overlaid (right). **j**, Clustered heatmap showing differential neighborhood distance analysis for clusters of the aging heart (old vs. young).

For the peripheral organs, markers for most of the expected cell types were identified, and no major deviations in size and location of each cluster were observed between the aging groups (**Extended Data Fig. 3a-e**). Nevertheless, comparing the expression for each annotated cluster between young and old mice returned a differentiated picture, with the kidney showing more than ten times the number of significant differentially expressed genes (DEGs) than the heart (**Fig. 1e**). Thereby, the peripheral organs each retained an almost disjunct aging-driven signature of dysregulation with notably varying effect sizes (**Fig. 1f, Extended Data Fig. 3f-j, Supplementary Table 3**). Interestingly, the peripheral gene expression clusters with the most DEGs were those known to harbor many immune cells (**Fig. 1g**). Yet, we found strong aging DEGs in each organ directly related to tissue function (**Fig. 1h**), such as genes of the myosin heavy chain in the heart (Myh7), members of the cytochrome P450 family in the kidney (Cyp2d12), or major urinary protein coding genes in the liver (Mup7). We wondered however, whether tissue-specific DEGs exhibited a spatially variable domain, independent of our supervised spot annotation. Thus, we determined the list of highly significant and reproducible spatially variable genes (SVGs) for each organ without making use of the spot cluster labeling (cf. Methods, **Extended Data Fig. 4a, b**, **Supplementary Table 4**). Similarly to the DEGs, the number of SVGs was organ-specific and we found that per tissue between 5% (spleen) and 24% (kidney) of the genes were both differentially expressed and spatially variable (**Extended Data Fig. 4c**). Still, except for liver and spleen, the number of SVGs clearly outweighed the number of aging DEGs. As one example, the prolactin hormone receptor (Prlr), a type I cytokine receptor, was induced upon aging specifically in the inner and outer stripe of the outer kidney medulla (**Fig. 1i**). Prolactin accumulation was previously reported as a sign of renal failure^14^. Another way to leverage the spatial resolution of the data is by investigating whether the local neighborhood, i.e. tissue microenvironment depends on the age (cf. Methods), since we found only minor changes in cluster proportion along the aging trajectory. Indeed, we again found tissue-specific neighborhood alterations, such as a progressive depletion of fibroblasts in the myocardium of the aging heart (**Fig. 1j**), or the loss of Kupffer cells in adjacency to periportal hepatocytes in the aging liver (**Extended Data Fig. 4d-g**). The immune system appeared to be implicated in all peripheral organs with increasing age, suggesting a more systemic relationship, especially considering the autocrine effects of parts of the dysregulated transcriptome such as the Prlr gene, which is typically also mediated through the central nervous system.

## Spatial aging signatures across the mouse brain

To better capture the structural and functional complexity of the murine brain, we decided to profile each sample at five different bregmata, using expert annotations from the Allen Mouse Brain Atlas for guidance (**Fig. 2a**). We observed well reproducible cluster gene signatures for each brain section (e.g. cortical layers I-Vi) with highly preserved marker expression between adjacent slides and used a previously established nomenclature to annotate the spot clusters^9^ (**Fig. 2b**, **Supplementary Table 5**). Similar to the peripheral organs, differences in observed cluster proportions were driven by the bregma and not by age. There was however, a trend for an overall decreasing clustering complexity for lower bregma (**Extended Data Fig. 5a-e**), due to the increasing area assigned to the midbrain structure. As expected, the different brain sections shared most of the DEGs due to similar constituting clusters, with a notable peak in Brain5, the only slice already covering parts of the cerebellum (**Fig. 2c, d**). Overall, the global effect sizes of aging DEGs in the brain were comparable to the peripheral organs, although much lower in their absolute number (**Extended Data Fig. 5f-j**, **Supplementary Table 6**). Interestingly, across all bregmata the number of SVGs clearly outweighed the number of DEGs by a factor of almost 100 (**Extended Data Fig. 6**, **Supplementary Table 7**). Still, the 17 aging DEGs shared between all five brain sections showed an enrichment of immune-system-related and disease-associated functions, such as C1qa-c and Trem2, while also exhibiting the strongest dysregulation in the white matter tract (**Fig. 2e**). Among those, 16 genes are also SVGs in the brain, demonstrating the need for transcriptomics in spatial dimensions to better understand the directions of aging-driven dysregulation. In particular, Egr1, Gfap and Trem2 are all known markers for neurons, astrocytes and microglia, respectively, triggering the question whether certain cell types specifically contribute on their own to the immune activation commonly observed during brain aging.

**Figure 2:**
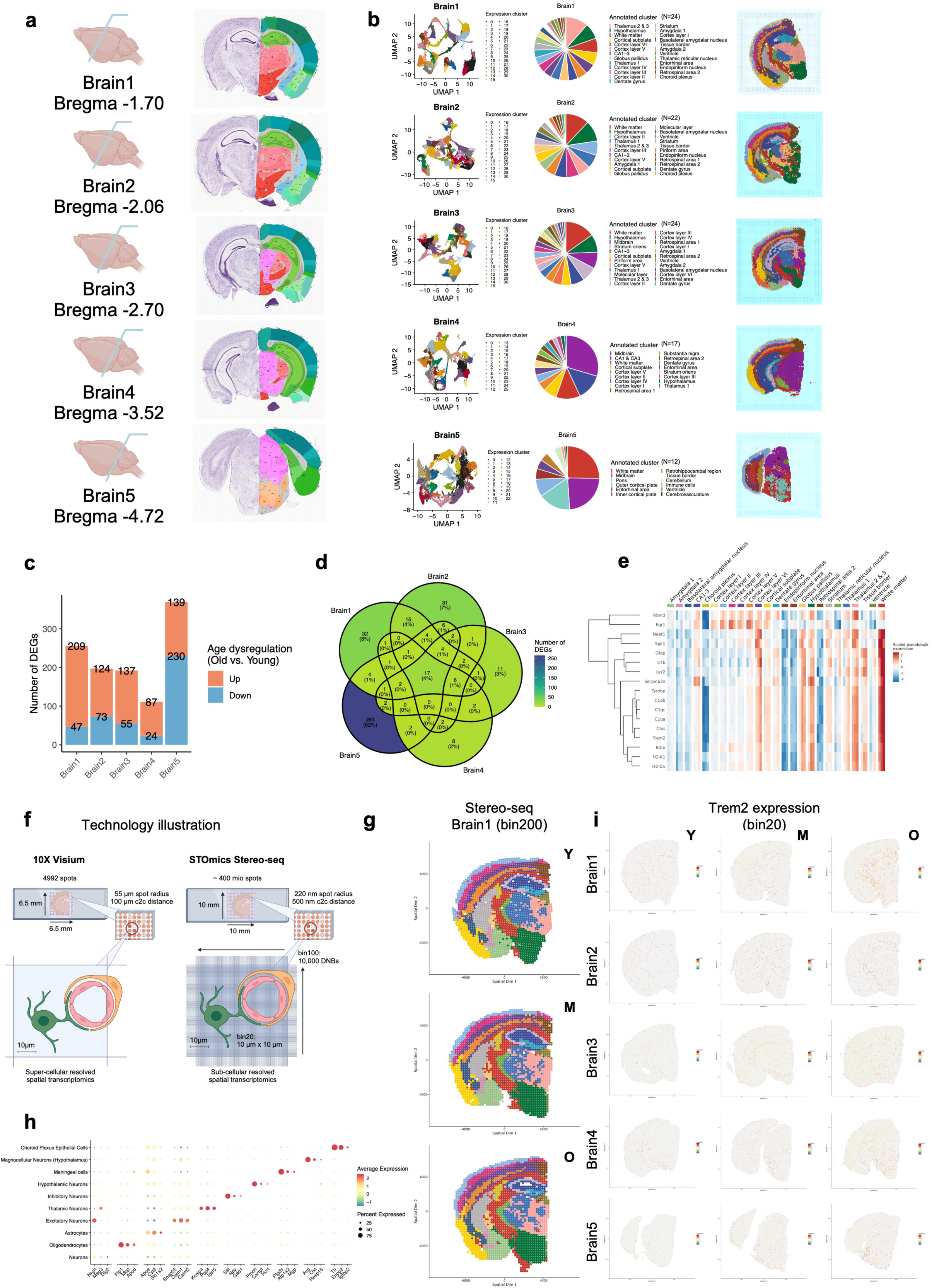
Spatial transcriptomics across the aging mouse brain. **a**, Visualization of the five Bregmata selected to study different regions of the aging brain. **b**, From left to right: Two-dimensional UMAP representation of colored spot clusters computationally integrated by brain slice (top to bottom), pie chart showing the proportion of annotated clusters across all brain samples, one representative annotated Visium sample with the spot cluster identities plotted over the H&E-stained tissue image. **c**, Number of differentially expressed genes per aging brain bregma (old vs. young) and direction of dysregulation. Total DEG counts were derived across all organ clusters, without removing duplicates. **d**, Five-dimensional Venn diagram comparing the brain DEG sets from (**c**). **e**, Heatmap showing scaled expression of the 17 aging DEGs (rows) shared between all five brain slices, using the Brain1 pseudobulk samples and spot clusters for visualization (columns). All genes except Rbm3 are also significant SVGs in at least one of the five brain bregmata. **f**, Sketch of 10x Visium and STOmics Stereo-seq examples comparing the features of both spatial transcriptomics technology platforms. **g**, Examples for binned (bin200) and annotated spot clusters of the aging brain (top to bottom; young, middle, old) at Bregma#1 sequenced with Stereo-seq. **h**, Dot plot showing the top 3 most significant marker genes per cell type annotated spot cluster using the Stereo-seq cellbin resolution of Brain1 samples. **i**, Normalized spatial expression of Trem2 across all 15 STOmics Stereo-seq brain samples using the near-cellular resolution bin20 (from left to right: young, middle, old; from top to bottom: Brain1-5). For visualization spot sizes were rescaled into the point interval [0.1, 1.5] according to their expression of Trem2.

To follow up on this observation, we validated our analysis of the super-cellular 10x Visium brain experiments with the sub-cellularly-resolved Stereo-seq^15^ technology developed by BGI STOmics, sparing one mouse brain from each age group (**Fig. 2f**). To evaluate concordance after quality control, we first aggregated the Stereo-seq spots into bins, artificially replicating the size of Visium spots. This allowed us to reproduce most of the gene expression clusters for each bregma and age but with slightly varying marker genes (**Fig. 2g, Extended Data Fig. 7, Supplementary Table 8**). All of the 17 shared brain aging genes from the Visium data set were successfully reported as DEG in at least four of the five matching brain layers in the Stereo-seq dataset, corroborating our findings (**Supplementary Table 9**). Next, we utilized the specific capabilities of Stereo-seq to perform an image-based cell binning of all spots near DNA fluorescence signals followed by cell type clustering and annotation. This enabled us to differentiate several types of neurons from common glia cells in the spatial domain (**Fig. 2h**). With our close-to-cell resolution (bin20), we confirmed the elevated expression of Trem2, a well-known microglia activation marker^16^, throughout the whole aged mouse brain (**Fig. 2i**). Taken together our findings further suggest a progressive dysregulation of the innate immune system that co-occurs with an increased spatially-dependent prevalence of major brain cell types in proinflammatory states.

Because normal brain function relies on numerous neurochemical pathways in different regions, presumable affected by aging through vascular-related oxidative stress, dysregulated glycolysis, or neurotransmitter dysfunction^17–19^, we prepared high-throughput spatial metabolomics experiments for the aging cohort using the other brain hemispheres for a set of tissue slices at Bregma 2.06 (cf. Methods). Each sample was then imaged in a 20 x 20 µm matrix grid to obtain per-pixel peaks. Obtaining four replicates per age group, we detected 2713 unique features across the 12 slices after computationally aligning the transformed MALDI timsTOF fleX mass readouts between samples, of which 97 peaks were reference-annotated to known metabolites (**Supplementary Table 10**). After a stringent quality filtering, 909 mass peak features remained for down-stream analysis. Among the strongest fold-changes (log(FC)=-9.7) between aged and young mouse brains we found N-Acetyl-L-aspartic Acid, located in the regions of white matter, striatium as well as the deeper cortical layers, even though statistical significance diminished after FDR correction (**Extended Data Fig. 8**). Thus, we argue that existing spatial metabolomics platforms are not yet sensitive enough to unbiasedly discover metabolome-wide changes in the context of brain aging, where often only subtle changes occur over larger time differences, but are better suited to perform targeted validation at cellular or sub-cellular levels^20^. The N-Acetyl-L-aspartic Acid molecule is a required metabolic factor for enabling lipid and myelin synthesis in oligodendrocytes^21^. We confirmed this finding in a comprehensive aging mouse brain atlas describing differential metabolomics in individually dissected anatomical regions, where N-Acetyl-L-aspartic Acid was also found be upregulated with aging, but specifically in the olfactory bulb^22^.

## Bulk and spatial transcriptomics of mouse organs in acute malaria

Severe infectious diseases (e.g. long COVID, tuberculosis, hepatitis, malaria) are associated with accelerated biological aging of the immune system in humans^23–25^. To dissect how organismal aging is (dis-)similar to systemic infection at the molecular level and which parts of the immune system activation are shared at the gene and pathway level, we repeated the 10x Visium experiments with a separate cohort of mice (cf. Methods). The infection cohort consisted of five young mice infected with the mouse parasite *Plasmodium berghei*, an animal model of cerebral malaria^26^ and five age- and gender-matched young controls (**Fig. 3a**). Because heart and spleen showed rather small effect sizes in the aging cohort, sections from the infection cohort were processed for three peripheral organs (liver, kidney and lung), in addition to two preselected brain regions at bregma –2.06 and bregma 3.89 (the olfactory bulb), the latter of which is the very first affected brain structure in this model^26^. After QC, we retained 42 high-quality capture areas with comparable spot statistics as mentioned above (**Fig. 3b**, **Extended Data Fig. 9**, **Supplementary Table 11**). The angles of histological slices were analogously chosen to best capture the tissue complexity, yielding a highly similar number, type and spatial distribution of spot clusters as for the aging cohort (**Fig. 3c, d**, **Supplementary Table 12**). However, here we noticed significant differences in the spot cluster proportions between the infected mice and controls. For example, the loss of blood vessels in kidney and liver parallel to a massive increase of immune cells in the liver and lung (**Extended Data Fig. 10a-e**). In line with this observation, the total number of DEGs in the infection cohort was much higher than for the aging cohort, with most being strongly upregulated after infection in a tissue-specific manner, meeting our expectations of an acute inflammation resembling a much more severe phenotype than aging (**Fig. 3e, f**, **Supplementary Table 13**). In particular the leukocyte, blood and lymphatic vessels as well as endothelial spot clusters exhibited the most DEGs and highest effect sizes across tissues (**Extended Data Fig. 10f-k**). In stark contrast to the aging cohort, the reproducible number of SVGs exceeded the number of DEGs exclusively in the brain, but with similar percentages of overlap per tissue, reaching from 7% in the olfactory bulb to 19% in the kidney (**Extended Data Fig. 11a-c**, **Supplementary Table 14**). The substantial expansion of connective tissue around the lung pulmonary veins together with a massive leukocyte invasion represented an intriguing phenotype alteration of the infected lung, which is immediately visible from the spatially annotated gene expression data alone (**Fig. 3g**). All three peripheral organs underwent remarkable changes in their spatial microenvironment, like the increased proximity between Kupffer and other immune cells in the infected liver, as revealed through a differential spot neighborhood analysis (**Extended Data Fig. 11d-f**).

**Figure 3:**
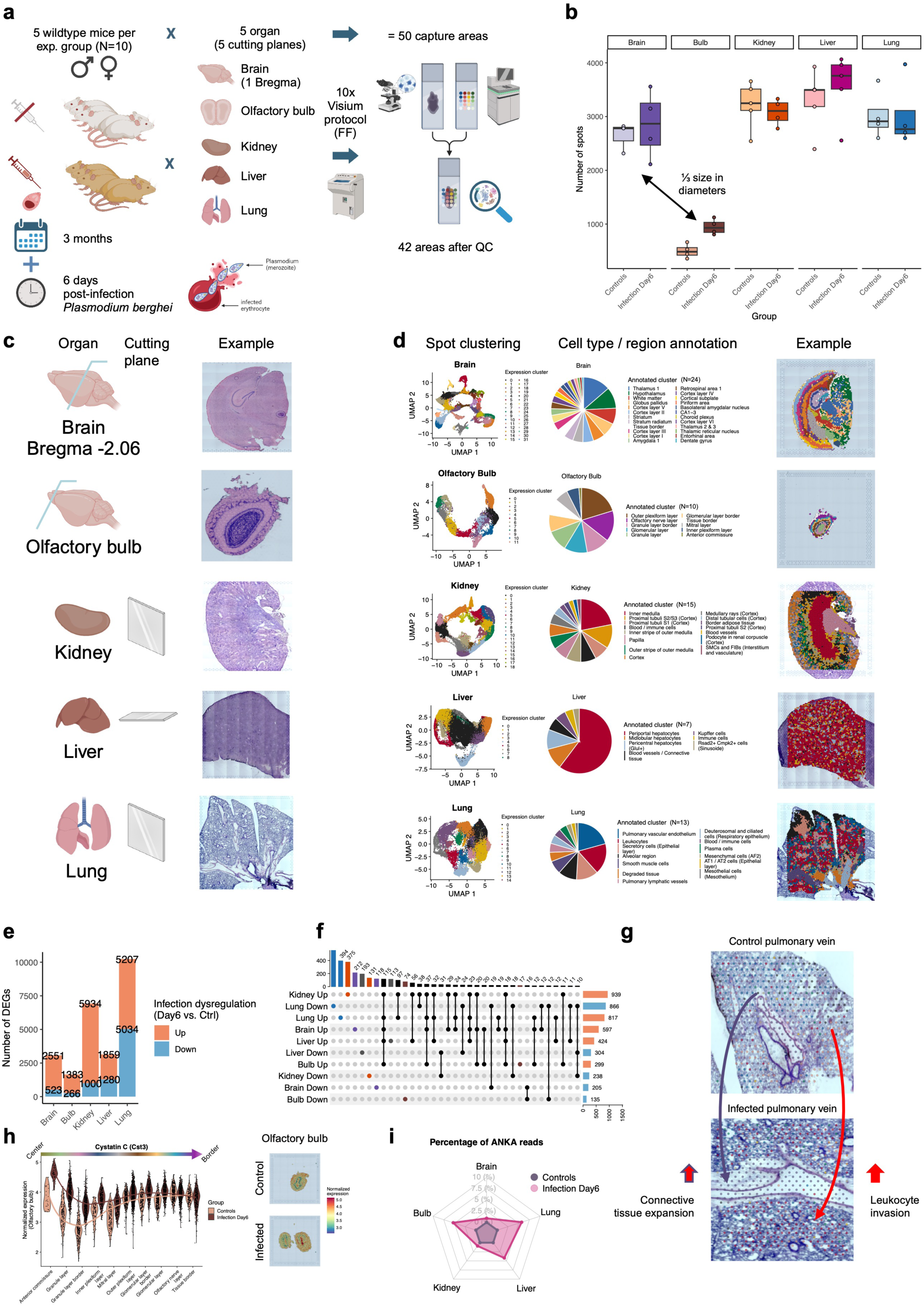
Spatial transcriptomics of the acute malaria-infected mouse. **a**, Illustration of the experimental workflow conducted for the malaria infection cohort. **b**, Distribution of the number of cleaned spots under tissue per organ and experimental group (controls, *P. berghei* infection). Average number of cleaned spots located under tissue varied between organs (Brain: 2760, Olfactory bulb: 722, Kidney: 3154, Liver: 3435, Lung: 3024), out of the 4992 spots possible per capture area. In general, the number of spots per sample was influenced by slight inter-individual variances in organ size. Colored boxes span the first to the third quartile with the line inside the box representing the median value. The whiskers show the minimum and maximum values or values up to 1.5-times the interquartile range below or above the first or third quartile if outliers are present (shown as separate, colored dots). **c**, Illustration of cut angle plane and one representative H&E-stained capture area per organ. **d**, From left to right: Two-dimensional UMAP representation of colored spot clusters computationally integrated by organ (top to bottom), pie chart showing the proportion of annotated clusters across all samples, one representative annotated Visium sample with the spot cluster identities plotted over the H&E-stained tissue image. **e**, Number of differentially expressed genes per organ (day 6 post *P. berghei* infection vs. controls) and direction of dysregulation. Total DEG counts were derived across all organ clusters, without removing duplicates. **f**, UpSet plot comparing the gene sets shown in (**e**). **g**, In-depth zoom into a lung pulmonary vein from a control mouse (top) and an infected mouse (bottom), demonstrating the expansion of connective tissue and leukocyte invasion caused by the malaria disease. Spots are colored according to cluster identity, in particular Leukocytes in red, Smooth muscle cells in purple and Blood / immune cells in light purple. **h**, Normalized expression of Cystatin C (Cst3) in all spot clusters of the olfactory bulb split by infection and controls with clusters ordered by their ascending distance to the center of the tissue (left). Trend lines were computed with loess regression of y ∼ log(x). Shown on the right is the normalized expression of Cst3 in the spatial domain of one representative control (top) and infected mouse (bottom). **i**, Percentage of reads per spot mapped to the *P. berghei* ANKA genome split by experimental group and organ.

Since *P. berghei* causes cerebral malaria and strongly affects the olfactory bulbs, we investigated the intersection of SVGs and DEGs more closely to understand the underlying mechanism. In general, many immune cell surface genes from the clusters of differentiation were significantly upregulated at specific locations in the infected olfactory bulbs, including Cd151, Cd164, Cd1d, Cd200, Cd24a, Cd44, Cd52, Cd55, Cd63, Cd74, Cd82, Cd9, and Cd99l2. Moreover, it appeared that antimicrobial factors such as Cystatin C (Cst3) were highly upregulated in the center of the infected olfactory bulb. This dysregulation pattern completely diminished toward the outer layers, thus showing an intriguing spatial expression pattern (**Fig. 3h**). Because *P. berghei* is a eukaryotic pathogen, we exploited the fact that both host and pathogen transcribe polyadenylated messenger-RNA, such that in principle both should be captured through the reverse poly-T tag in 10x Visium 3’ gene expression libraries. Indeed, we were able to map a certain percentage of high-quality reads from each infected sample against the *P. berghei* ANKA genome, with olfactory bulb, liver, and lung exhibiting the highest percentage of cells with pathogenic reads (**Fig. 3i**). We were thus able to narrow down the position of individual parasites within the tissue by following the distribution of pathogenic reads into the different spot expression clusters (**Extended Data Fig. 12**). Our analysis further revealed that they preferentially reside in the outer layers of the olfactory bulb, rendering the expression of Cst3 reciprocal to the distance between center and parasite location. We then identified the spots in the olfactory bulb clearly enriched for parasite reads and compared the gene expression of each such center spot with its six direct neighbors in the hexagonal spot array on 10x Visium capture areas. Neighbors of a parasite-enriched center spot exhibited a significant activation of the cytokine CC-chemokine ligand 5 (Ccl5) as well as the cluster of differentiation 74 (Cd74), indicating an acute T cell response around the pathogens. Such a severe immune system activation is a necessary means to fight infections.

Alteration of mRNA expression is modulated by several independent mechanisms, but partially explained by post-transcriptional gene regulation. We previously showed that both tissue-specific and global microRNA (miRNA) regulators are either activated or deactivated upon aging in the mouse^4^. We therefore wondered whether such a regulatory interplay also explains the spatial immune activation observed during infection and performed small non-coding RNA sequencing for the infection cohort (cf. Materials and Methods, **Supplementary Table 15**). We revealed that variability of miRNA expression is strongly driven by tissue origin and not by technical bias (**Extended Data Fig. 13a, b**). In total, 542 miRNAs were well expressed in at least one tissue and within one of the experimental groups (**Extended Data Fig. 13c**). The observed expression was highly correlated among samples from the same tissue (**Extended Data Fig. 13d**). Surprisingly, none as well as only one of the miRNAs remained significantly differentially expressed for brain and and bulb (mmu-miR-3473b, up), respectively (**Extended Data Fig. 13e, f**). For the peripheral organs however, we obtained 46 up-regulated miRNAs in kidney, 188 up-regulated and 12 down-regulated miRNAs in liver and again none for lung (**Extended Data Fig. 13g-i**), suggesting once more a tissue-specific gene regulatory mechanism to be involved in the systemic immune response to malaria disease. Looking into their potential function, we found the target gene Cd28 (adj. p=0.0021318), an essential factor for T-cell proliferation and cytokine production and the GO terms “positive regulation of inflammatory response to antigenic stimulus” (adj. p=0.0029171), and “positive regulation of peptidyl-serine phosphorylation” (adj. p=0.0029986). For the up-regulated liver miRNAs we found an enrichment of miRNA targets controlling “angiogenesis” (adj. p=4.79e-7) and “macrophage differentiation” (adj. p = 1.58e-6). All of these observations are in line with our hypothesis that miRNAs mediate peripheral immunity specifically and locally, triggering distant cellular interaction in the central nervous system and further trans-dysregulation of brain or other spatially restricted mRNAs. Still, it raises questions as to whether such an inflammatory process causes permanent organ damage and how in contrast the aging-induced immune activation may initially cause more attenuated but, in the end, similar disease phenotypes.

## Shared spatial dysregulation in aging and systemic infection

To better understand similarities and differences between organismal aging and systemic infection using our high-resolution spatiotemporal transcriptomics readouts, we correlated the effect sizes between aging and infection cohorts for the same tissues and expression clusters. Here, the brain showed a remarkable resemblance between aging and infection, with the complement component C4b being one of the top upregulated genes (**Fig. 4a**). Similar trends were observed in the peripheral organs (**Supplementary Table 16**). C4b is part of the early complement pathway, specifically upregulated in the white matter tract of the aging brain, also upregulated in the infected brain but beyond the white matter area, thus showing a different spatial presence (**Fig. 4b**).

**Figure 4:**
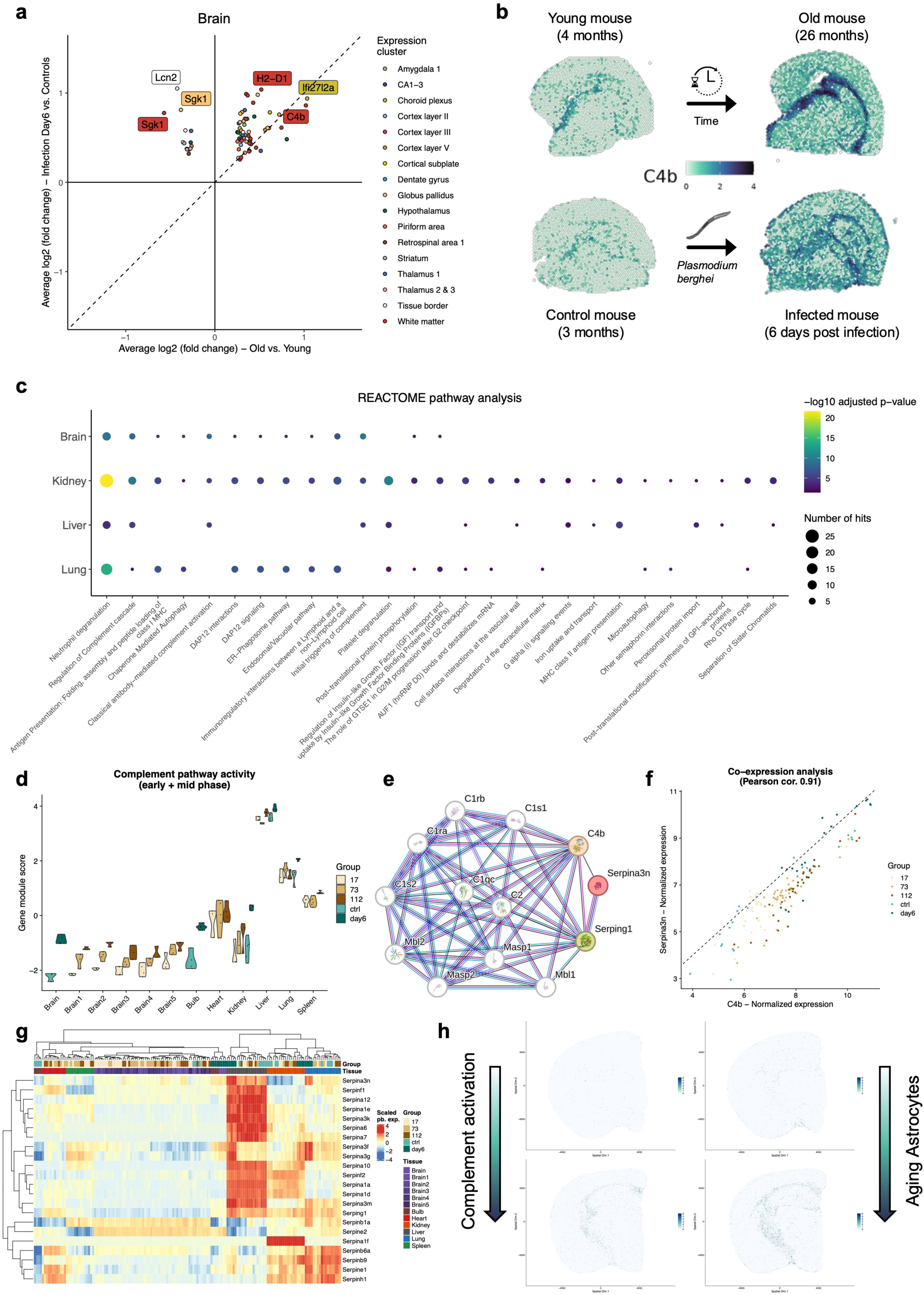
Molecular niches shared between aging and systemic infection. **a**, Scatter plot showing the average log2-scaled fold-change between aging and infection for all brain DEGs matched and colored by spot cluster. **b**, Normalized spatial expression of C4b in four representative brain samples (Bregma: -2.06), two from aging (top left: young, top right: old) and infection (bottom left: control, bottom right: infected) cohort. **c**, Dot plot showing adjusted and log-scaled Hypergeometric test p-values and the number of gene hits for the enriched categories from the Reactome pathway database across the four organs shared between aging and infection cohort. For each row of results a different list of genes was used as input (cf. Methods). **d**, Relative (scored) complement pathway activity in pseudobulk samples across all aging and infection cohort organs, split and colored by the five experimental groups (young, middle, old, healthy controls, infected). **e**, STRING network for Serpina3n in Mus musculus after performing one level of network expansion and removing edges from text mining and gene fusion. Edges are colored according to the type of association or interaction: curated databases (light blue), experientially determined (purple), gene neighborhood (green), gene co-occurrence (dark blue), co-expression (black), or protein homology (light purple). Nodes are colored according to their shell of interactions towards Serpina3n. **f**, Scatter plot showing normalized pseudobulk expression of C4b (x-axis) against Serpina3n (y-axis) across all samples and colored by the five experimental groups (young, middle, old, healthy controls, infected). **g**, Row and column clustered heatmap showing the scaled pseudobulk expression for all members of the serine protease inhibitor (Serpin*) gene family. **h**, Demonstration of spatially connected (sub-)cellular activity of the complement pathway (C4b+) and activated Astrocytes (Gfap+) in the aging brain (Bregma -2.06). Normalized expression values from one young representative on the top and one old representative on the bottom originating from the Stereo-seq samples at bin20 resolution are displayed.

We then performed an over-representation analysis for all the cluster-specific DEGs shared between infection and aging cohorts, and obtained “Neutrophil degranulation” and “Regulation of complement cascade” as top significantly enriched categories for brain, kidney, liver and lung (**Fig. 4c**, **Supplementary Table 16**). To validate this finding, we first created a gene signature feature for the complement pathway to measure its activity through gene module scores. Indeed, the spatially restricted activity of the early complement pathway was a common molecular feature of dysregulation between organismal aging and systemic infection across multiple organs (**Fig. 4d**). Since the enrichment analysis yielded the “Regulation of complement cascade”, we next wondered which gene regulatory components known from the literature are centered around C4b, revealing with STRING^27^ a dense network of other complement genes but also two serine protease inhibitors (Serpina3n, Serping1) (**Fig. 4e**). Surprisingly, Serpina3n was one of the top dysregulated genes in the aging brain and also upregulated in the infected brain. Leveraging aggregated pseudobulk expression, we confirmed the almost perfect intra-sample correlation between C4b and Serpina3n across both cohorts and all organs in common (**Fig. 4f**). These findings suggested a central gene regulatory role between serin proteases involved in complement activation and serine protease inhibitors, mediating immune responses in two physiologically different processes that each depend on extrinsic and intrinsic stimuli. We then looked into the whole family of Serpin genes and discovered a tissue-specific dysregulation of its members in aging and infection (**Fig. 4g**). Subsequently, we made use of our sub-cellularly resolved Stereo-seq data sets to closely compare the complement activity against the activity of reactive astrocytes that drive parts of the brain aging signature and obtained a high degree of spatial correlation (**Fig. 4h**). Utilizing our respective marker-based cell bin annotations, we deconvolved the spatial pattern of correlated C4b and Serpina3n expression to primarily originate from both from Astrocytes and Oligodendrocytes. We confirmed this for the cell-marker enriched spots in the aging and infected brain Visium samples (**Extended Data Fig. 14a, b**). Moreover, we compared our findings with a previous study^4^ describing in-depth cross-organ bulk miRNA and mRNA expression in aged mice, finding indeed multiple members of the Serpin gene family (Serpina1f, -3f, -3g, -3k) as anticorrelated targets of aging-associated miRNAs in several tissues. Finally, we externally validated a significant aging-associated upregulation of C4b and Serpina3n on the protein level in the mouse brain cortex and hippocampus, using a previously published proteomics data set^28^ (**Extended Data Fig. 14c**).

In sum, our findings indicated a cross-organ gene regulatory mechanism as part of the immune system, being centered around serine proteases (**Extended Data Fig. 15**). This dysregulated pathway appeared to be orchestrated in each organ in a cell type-specific and spatiotemporal manner and translated from the transcriptional level into differential protein abundances, increasing the level of confidence for functional relevance.

## Discussion

Understanding the heterogeneous and physiological consequences of mammalian aging is a challenging task. Our large-scale spatiotemporal cross-organ study provides new molecular insights into the underlying mechanisms. Several spatially and temporally resolved genomic data sets on the mammalian brain^9,29–33^, olfactory bulb^34–36^, heart^37–46^, kidney^47–51^, liver^52–54^, lung^55–61^, and spleen^62–65^ have been published, each describing their molecular architecture in health and disease. However, most of them lacked a sample size sufficient to compare multiple experimental groups and organs in parallel, especially lacking from the same donor^66^. Large sequencing studies are inherently challenging to perform and thus carry certain risks of producing unwanted batch effects^67^. We tackled these challenges by utilizing two separate breeds of wildtype mice, one from the United States for the aging cohort and one from Germany for the infection cohort, while ensuring a minimum number of three replicates per experimental group. The aging cohort was processed first and fully completed before continuing with the infection cohort to prevent potential sample cross-contamination.

Our findings suggest shared physiological processes between aging and infectious diseases, at least in malaria, that are progressively dysregulated and are centered around complement activation and serine protease inhibitors. C4b and Serpina3n were previously reported to be upregulated in the subventricular zone of the aged mouse brain and as being indicative of a reactive type of oligodendrocyte^68^. Yet, Hajdarovic et al. recently reported an activation of C4b in reactive Astrocytes from the female hypothalamus and hippocampus, suggesting a more general, sex-specific role of glia cell types in mediating a neuroinflammatory or autoimmune phenotype^6,69^. Moreover, Allen et al. narrowed down C4b upregulation in the aged mouse brain in both astrocytes and oligodendrocytes that was further escalated by LPS treatment ^7^, also corroborating our findings. Others reasoned that such a phenotype could solely arise from the body-wide accumulation of senescent cells or an organ-dependent dysregulation of the circadian clock, two other hallmarks of aging^12,70^. In humans, C3 and C4 levels were reported to be anti-correlated with centenarian longevity^71^. It therefore remains intriguing, having perceived an almost perfect correlation between C4b and Serpina3n across the aging and infection cohorts, to raise further questions about the mutual gene regulation of complement proteases and protease inhibitors, and conceivable downstream effects. Proteases play an important role in innate immune defense mechanisms, but need to be tightly regulated in order to prevent autoimmunity and self-attack, and thus have already been brought in connection with age-related decline and neurodegeneration^72^. Within the complement system, C4 is cleaved through the serin protease activity of C1 in the classical pathway and through MBL-associated serine proteases (MASPs) within the lectin pathway^73^. Both proteases can be inhibited by Serping1, a serin protease inhibitor^74^, which is upregulated in the aging mouse heart and kidney as well as during malaria infection of kidney, brain, olfactory bulb, liver and lung. We were able to confirm for cortex and hippocampus that indeed the upregulation of C4b and Serpina3n RNA transcripts results in increased protein abundance. Besides, Serpina3n is another dysregulated serine protease inhibitor of chymotrypsin, like Serping1, but is mostly known for the regulation of Cathepsin G^75^. After glycosylation, Serpina3n can also bind double-stranded DNA in the nucleus and cause chromatin condensation^76^. Whether the observed correlation of C4b and Serpina3n is explainable through its regulatory capacity within the nucleus can only be determined in further proteomics studies with functional validation assays. In general, looking beyond the complement system, the distribution of lymphoid and myeloid cells in aging tissues, as well as their contribution to sterile inflammation during the aging process is being increasingly recognized as highly complex and tissue-dependent^77^. Our data as well as previous reports^78^ point toward a cross-communication of native tissue parenchymal cells with lymphoid cells that is implicated in the different tissue-specific aging niches, for instance the white matter compartment in the brain or the lung alveoli. A more targeted and deeper approach using single-cell or spot-free spatial *in situ* mapping techniques to quantify the contribution of myeloid and lymphoid cells toward the development of shared aging- and infection-driven niches and how this is mediated through the peripheral systems, is needed. And even though the high-throughput analysis of the aging brain metabolome has neither directly confirmed nor invalidated our findings on C4b, we have detected almost 1000 yet uncharacterized mass feature peaks exhibiting strong fold-changes in spatially restricted zones. Further research is needed to decode their individual structure and function, leaving high potential for a comprehensive understanding of the metabolome and its role in brain aging.

Moreover, it remains to show whether the effects described here occur in the same or similar way in human tissue, given the expression of cell markers obtained with single-cell experiments was already shown to be partially discordant between species^79^. In fact, an evolutionary discrepancy determines in any case the baseline chances of success to carry over our findings into new therapeutic strategies targeting aging and infectious diseases. With the establishment of improved spatial transcriptomics platforms capable of resolving RNA expression at the sub-cellular level, we will be able to better narrow down the origins of individual RNA classes and molecules, for instance whether they are enriched in the nucleus or the cytosol. This will help to resolve missing details about the so far hidden cell to cell crosstalk involved in complement biology as just recently suggested^80^. Our findings position individual members of the complement system and protease inhibitor families as potential broad therapeutic targets in progressive aging and infectious disease research. Such targets are desperately needed to tackle pathogenicity, as for example cerebral malaria is still lethal for approximately 20% of affected patients, with many survivors being affected by prolonged neurocognitive impairment after primary therapy^81^. We therefore reason that a complement activation-based pathomechanism plays an important role in autoimmunity-related tissue damage that in turn contributes to diverse aging- and infectious disease-related phenotypes.

Beyond aging research, single-cell and spatial transcriptomics are pivotal techniques to study the host-pathogen interface and reveal specific biological mechanisms such as cell invasion or clearance^82,83^. While most bacteria and virus specific protocols still have to be developed in order to dual-detect both host and pathogen from the same sequencing sample, multicellular parasite species can be directly sequenced due to the same type of polyadenylated messenger-RNA as in the host^84^. This compatibility was already broadly used to describe the organismal development of *P. berghei*, mechanisms behind its transmission as well as specific effects on gene expression induced in individual organs and cell types^65,85–91^.

We initially combined the existing literature knowledge on individual tissues and cell types as well as several independent single-cell and spatial studies to carefully annotate the spot gene expression clusters needed for the downstream analysis. Unfortunately, there exists no common data annotation standard in the community, even though a few interesting nomenclatures and solutions to single-cell reference atlases were already proposed^92–94^. Our data set is specifically designed to foster spatiotemporal driven research into specific organs and cell types that are of particular interest in the aging and infection research communities, but also beyond to serve as a reference data set for any other biomedical domain. However, analyzing almost 200 Visium samples in an integrative approach brings also significant computational challenges. Many tools are not ready to scale up to large samples sizes, requiring extensive compute infrastructure and workflow optimizations. Therefore, we anticipate that our data set is also beneficial to the computational methods community, fostering the development of novel and scalable data analysis methods in the spatial transcriptomics domain.

## Methods

### Mouse tissue processing and screening

For the aging cohort, male C57BL/6JN mice were shipped from the National Institute on Aging colony (Charles River) to the Stanford ChEM-H animal facility, where they were housed at 12/12h light/dark cycle in cages of 2-3 mice. Water and food were provided ad libitum. The aging cohort consisted of 5 mice per age group (age groups: 17, 73 and 112 weeks; 3.91, 16.8 and 25.78 months). Tissues from brain, heart, kidney, liver, lung and spleen were collected after anaesthetization with 2.5% v/v Avertin and transcardial perfusion with 20 mL cold PBS and flash frozen in isopentane cooled with liquid nitrogen, all between 10:15 AM-12:00 PM on a single day. For brain samples, the right and left hemisphere were separated before freezing and stored separately. Samples were stored at -80°C until embedding in OCT. Samples were embedded for sectioning in precooled OCT on dry ice. All animal care and procedures complied with the Animal Welfare Act and were in accordance with institutional guidelines and approved by the institutional administrative panel of laboratory animal care at Stanford University.

For the infection cohort, experimental procedures were conducted as described previously^95^. In brief, C57BL/6 mice were bred at the animal facility and *P. berghei* ANKA parasites were obtained from the blood of sporozoite-infected C57BL/6 mice provided by the Parasitology Section of the Bernhard Nocht Institute. Mice in the control and experimental groups were age-matched (6 to 8 weeks old) and sex-matched (male & female). For the experimental group, C57BL/6 mice were infected i.p. with 1×10^5^ parasitized red blood cells. Positive parasitemia was determined in Giemsa-stained blood smears from tail blood. Body weights were determined at different time points. All required organs were collected on two consecutive days, from 9:30 AM (Ctrl.) and 10:30 AM (day 6 post-infection) until 12 PM. To prevent lung deflation, the tracheae were sutured, and the lungs were inflated with a 1:1 mixture of PBS and OCT. The small lung lobe was separated and frozen separately from the residual lung tissue. Brains were frozen without separating the hemispheres. All tissue samples were simultaneously frozen while embedded in OCT, in an isopentane bath cooled with liquid nitrogen and kept stored at -80°C. All animal care and procedures complied with institutional guidelines and were approved by the administrative panel of laboratory animal care at the Bernhard Nocht Institute under the admission TVA N118/2020, as well as the Hamburg Animal Welfare agency.

Cryosectioning and section mounting were performed according to the manufacturer’s protocol, in a Leica cryostat (LEICA CM3050 S) set to -15°C with a slice thickness of 16 µm (except for the spleen of the infection cohort; 10 µm). For the aging cohort, five brain sections were mounted using visual landmarks and the Allen Mouse Brain Atlas, resulting in sections from bregma –1.7, -2.06, -2.7, -3.52 and -4.72. For the infection cohort, two brain sections were mounted, namely from bregma –2.06 and bregma 3.89 (Olfactory bulb). Sections were mounted on designated capture areas and stored at -80°C until further processing. After tissue mounting on the slide, an additional 10-15 sections were collected for RNA Quality assessment. All samples used in this study had a RIN value between 4.4 and 9.0 (**Supplementary Table 1, 11**).

### Spatial gene expression assay (10x Visium)

Following the Tissue Optimization protocol, we determined the permeabilization times to 18 minutes for the brain, 24 minutes for the olfactory bulb, 15 minutes for heart, 15 minutes for kidney, 16 minutes for liver, 20 minutes for lung, and 30 minutes for the spleen samples using the Tissue Optimization (TO) Slide & Reagent Kit (10x Genomics, PN-1000184).

We followed the standard protocol for fresh frozen tissues (compare document CG000239 - Rev F) using the Visium Spatial Gene Expression (GEX) Slide & Reagent Kits with recommended reagents (10x Genomics, PN-1000193). In brief, after H&E staining and imaging (10x magnification, Axio Imager KMAT - ZEISS), tissues were permeabilized with the previously defined permeabilization times. After reverse transcription on the slide and denaturation, second strand synthesis is performed. For further processing, cDNA is released via denaturation. To determine cycle number for amplification the KAPA SYBER FAST qPCR Master Mix Kit (KAPA Biosystems) is used in a qPCR. All samples in this study were amplified using 14 – 19 PCR cycles. After bead purification using SPRIselect beads (Beckman Coulter, Product No. B233318), cDNA is quantified via 2100 Bioanalyzer using the Bioanalyzer High Sensitivity DNA kit (Agilent, cat.no.:5067-4626).

The library construction was either performed manually or using the MGISP-100 liquid handling system according to the 10x Visium user guide. In short, the following steps were performed: fragmentation, end repair, A-tailing, double sided size selection, adaptor ligation, post ligation cleanup, sample index PCR, double sided size selection; followed by post library construction QC performed using the Bioanalyzer High Sensitivity DNA kit (Agilent, cat.no.:5067-4626) and KAPA Library Quantification Kit (Roche, KK4824).

Libraries were pooled in an equimolar fashion with up to 15 samples in one library, multiplexing only samples that originated from the same organ whenever possible. All libraries were sequenced using NovaSeq 6000 (Illumina).

### Spatial gene expression assay (STOmics Stereo-seq)

One brain sample of each age was processed at the MGI Tech Co., Ltd. (Riga, Latvia) using the STOmics Stereo-seq Transcriptomics T Kit (MGI). For each brain, five 10µm sections (Bregma: –1.7, -2.06, -2.7, -3.52 and -4.72) identical to the Visium cohort were collected and loaded onto the Stereo-seq Chip T-Slide according to manufactureŕs protocol. In short, after fixation of the tissue using ice-cold methanol, cell nuclei were stained using the ssDNA Qubit dye (Thermo Fisher) and imaged in 10x resolution in FITC channel. Subsequently, brain tissue was permeabilized for 12 min at 37°C, releasing mRNAs from the cells onto the Chip T-Slide. Reverse transcription of mRNAs was conducted for 3 hours at 42°C generating cDNA with sequence barcodes indicating their unique location on the T-Slide. Afterwards, the tissue was removed using tissue removal buffer for 10 min at 55°C and cDNA was collected from the chip o/n at 55°C. CDNA was then purified, amplified and size distribution was checked using High Sensitivity DNA Bioanalyser Kit (Agilent). Library Preparation was performed using Stereo-seq Library Preparation Kit (MGI) using 30ng of cDNA and 13 cycles of amplification following the manufactureŕs protocol. Final quality check of the size distribution of the library was performed using High Sensitivity DNA Bioanalyzer Kit. Libraries were sequenced on DNBSEQ-Tx sequencer (MGI).

### Small non-coding RNA profiling

Total RNA was isolated from 5-10 10µm tissue sections using the miRNeasy Mini Kit (Qiagen, Hilden, Germany) according to the manufacturers protocol. RNA concentration and integrity was determined using Nanodrop (ThermoFisher Scientific, Waltham, MA, USA) and RNA 6000 Nano Bioanalyzer Kit (Agilent, Santa Clara, CA, USA), respectively. Libraries for miRNA expression profiling were generated with the MGIEasy Small RNA Library Prep Kit (MGI Tech, Shenzhen, China) according to manufacturer’s recommendations with 100ng totalRNA starting material. Sequencing was performed on an DNBSEQ-G400RS instrument by the Sequencing Unit of the Core Facility Molecular Single Cell and Particle Analysis of Saarland University using the 50bp single end sequencing strategy.

### Image and data preprocessing

We stitched the image series from the Axio Imager KMAT brightfield microscope using the Zeiss ZEN software (v3.6) and cropped and rotated the resulting high-resolution TIFF files using Fiji. Individual sample images were color corrected where needed to match the other images. In the next step, we ran Spaceranger (v2.1) with the demultiplexed FASTQ files and the corresponding TIFF images. In brief, Spaceranger aligns the reads to a reference genome (refdata-gex-mm10-2020-A), detects tissue under spots and the fiducial frame in the image, to create a spot x UMI count matrix. For the infection cohort, we additionally built a custom Spaceranger reference genome combining mouse (GRCm39) and *Plasmodium berghei* - ANKA genome (PBANKA01). For two samples the alignment of the fiducial frame was manually corrected after the automatic detection had failed initially.

In case of the STOmics Stereo-seq samples, images were obtained and stitched with the Zeiss Moticam Pro 500i. Combined QC and primary analysis of sequencing and image data was performed using the STOmics analysis workflow (SAW) v1.0 to obtain quantification matrices for the mouse genome (GRCm38). The SAW pipeline was also used to accumulate DNB spot gene expression counts into the specific area and cell binnings.

### Bioinformatics and quality control

Based on the Spaceranger reports, we either excluded a sample completely from further analysis, or using the 10x Loupe browser tool (v6.5.0) manually selected small spot regions to exclude from downstream analysis where necessary (e.g. removing corners with uneven permeabilization), and further applied the following criteria-based spot filtering on each sample, to eventually obtain a high-quality data set. To this end, we inspected the variable distributions for the number of UMIs, the number of read counts, mitochondrial content, as well as spot complexity (=log10(Number of features) / log10(Number of counts)) for each tissue separately, and applied the following filtering criteria on the filtered spot count matrix from Spaceranger after loading into Seurat objects: (*Maximum percentage of mitochondrial read content; Minimum number of UMIs; Maximum number of UMIs; Minimum number of detected features; Maximum number of detected features; Minimum spot complexity*)

Brain: (35; 500; 70,000; 300; 10,000; 0.75), Bulb: (35; 500; 70,000; 300; 10,000; 0.75),

Heart: (60; 1000; 25,000; 300; 5000; 0.75), Kidney: (30; 1000; 50,000; 300; 8000; 0.75)

Liver: (15; 1000; 40,000; 500; 8000; 0.75), Lung: (10; 1000; 30,000; 300; 8000; 0.75)

Spleen: (10; 500; 35,000; 300; 8000; 0.75).

We then rotated the images of the remaining samples to best match the desired orientation for each tissue. We then followed the default Seurat (v4.4) workflow to preprocess each sample by applying *NormalizeData*, *FindVariableFeatures* with variance stabilizing transformation for 2000 features, followed by *SCTransform* (v1), regressing out the percentage of mitochondrial reads and subsetting to 3000 variable features. Subsequently, we performed *RunPCA* on the SCT assay, *FindNeighbors* on up to 30 PCs, *FindClusters* with leiden clustering followed by *RunUMAP* with up to 30 dimensions. Using the cleaned list of Visium samples, we created one Seurat object for each tissue using the SCT-based integration workflow as follows: set the default assay to SCT, run *SelectIntegrationFeatures* with 3000 features, then *PrepSCTIntegration*, and *FindIntegrationAnchors* followed by *IntegrateData*, each with “SCT” set as normalization method and up to 30 dimensions. Next, we performed the standard dimension reduction workflow consisting of *RunPCA*, *RunUMAP*, *FindNeighbors*, and *FindClusters* again with up to 30 dimensions and fixed random seeds.

For the STOmics Stereo-seq samples, we first converted the outputs of the Stereo-Seq Analysis Workflow (SAW) pipeline^96^ (v1.0) into Seurat objects for both bin200 (200×200 Stereo-seq spots) and cell bins. This was accomplished using the *io.stereo_to_anndata* function in the Stereopy package^97^ (v1.1.0), followed by applying the *h5ad2rds.R* script provided by BGI. Cell bins were obtained using image-based cell segmentation of the corresponding ssDNA-stained images. Filtering was performed using the following criteria: (*Maximum percentage of mitochondrial read content; Minimum number of UMIs; Maximum number of UMIs; Minimum number of detected features; Maximum number of detected features; Minimum spot complexity*)

Brain (bin200): (35; 500; 100,000; 300; 10,000; 0.75), Brain (cell bin): (35; 100; 10,000; 100; 3,000; 0.75).

In line with the Visium 10x samples we applied a default Seurat preprocessing workflow for each sample. Using the cleaned list of STOmics Stereo-Seq samples, we performed the Harmony-based integration workflow as follows: set the default assay to SCT, run *SelectIntegrationFeatures* with 5000 features, merge the sample list into one Seurat sample, run *RunPCA* with 50 components, run *RunHarmony* on SCT assay and the PCA reduction using *Sample_ID* and *Age* as latent variables, run JoinLayers on the RNA assay. Then, we performed the sample dimension reduction workflow as in the Visium samples with up to 50 dimensions.

### Spot clustering annotation

For the Visium aging cohort we employed tissue-specific clustering resolutions to match their individual complexity as follows: Brain1 0.8, Brain2 0.7, Brain3, 0.6, Brain4 0.5, Brain5 0.5, Heart 0.5, Kidney 0.5, Liver 0.2, Lung 0.5, and Spleen 0.6. For the Visium infection cohort, we set the clustering resolution for Brain to 1, Olfactory bulb to 0.6, Kidney to 0.5, Liver to 0.2 and Lung to 0.5, aiming to reproduce about the same number and type of gene expression clusters as for the aging cohort. In the next step, we computed the enriched cluster marker genes for each tissue object in a one vs. all approach by using *FindAllMarkers* on the normalized data slot of the spatial assay in combination with MAST^98^, setting the minimal percentage of positive cells to 0.1, requiring a minimal alpha to 0.01 and log fold-change of 0.1 for Heart, Kidney, Lung, Spleen as well as 0.25 for brain and using the sample identity of each spot as latent variable for gene regression. We then applied a hierarchical approach of annotating the clusters using the top-enriched genes, trying to resolve at best the most-enriched cell type, if possible, through the expression of clear marker genes, and annotating the spatial compartment based on histological knowledge and functional properties instead. We made combined use of transcriptomics literature on each organ, in particular single-cell and single-nucleus reference data, reference atlases such as the Allen Mouse Brain atlas, as well as an in-depth analysis of the stained tissue slices, to manually annotate all clusters with a nomenclature used previously. Thereby, clusters having no distinctive but unclear markers but showed shared marker expression overall to other clusters, were merged together.

For the STOmics Stereo-seq samples with bin 200 (200×200 DNB bins), we employed tissue-specific clustering resolution aiming to approximately reproduce the number of clusters from the Visium 10x aging cohort at each bregma as follows: Brain1 3.2, Brain2 3, Brain3 3, Brain4 2.5, Brain5 2.5. For stereo-seq samples with single-cell resolution we utilized a clustering resolution of 0.5 across all Bregmata. In the next step, we computed the enriched cluster marker genes for each tissue object in a one vs. all approach by using *FindAllMarkers* on the normalized data slot of the spatial assay in combination with MAST, setting the minimal percentage of positive cells to 0.1, requiring a minimal alpha of 0.01 and fold-change of 0.25 for all samples and using the sample identity of each spot as latent variable for gene regression. This was accomplished equivalently for both bin200 and cell bins. For bin200, we performed annotation by first calculating the marker overlap of each cluster with every cluster from the corresponding Visium 10x samples. Here, we always considered the top 50 markers selected by lowest adjusted p-value. To annotate a cluster, we then identified Visium clusters with marker overlap and additionally took the anatomical location, via the Allen Mouse Brain Atlas of the corresponding spots, into account. In most cases, this allowed us to pinpoint one annotation for each cluster. In cases where multiple annotations we plausible, we split the clusters based on spatial location and anatomical information. Clusters annotated with the same label were merged together. For cell bins we aimed for a high-level cell type annotation matching the limited counts per cell of the stereo-seq cell segmentation. We utilized the current literature on single-cell and single-nucleus datasets as reference for our annotation together with the anatomical location of the clusters. Spot clusters with similar markers and anatomical location were then merged together.

### Differential gene expression analysis

To compare the gene expression program of each tissue cluster between old and young mice as well as infected and control mice, we performed a DEG analysis using the *FindMarkers* function of Seurat in combination with MAST. Thereby, we used the normalized data slot of the spatial assay, requiring a minimal fold-change of 0.25, minimal percentage expressed of 0.1, as well as an alpha level of 0.01, while again using the sample identity of each spot as latent variable for gene regression. A gene was considered as significantly dysregulated for an organ if it returned as significant in at least one of the spot clusters.

### Spatially variable gene analysis

We applied SPARK-X^99,100^ (v1.1.1) to each Spaceranger-filtered and preprocessed but uncleaned sample matrix to detect the spatially variable genes in each tissue sample, independent of our clustering and annotation approach and thus to prevent any artifacts arising from our cleaning procedures. To this end, we used the SCT normalized counts as input to SPARK-X and selected the mixture projection kernel. We then only consider genes as spatially variable, if they are significant in the *mtest* slot to an alpha level of 0.01. For the comparison to the tissue cluster DEGs we further filtered the SVGs by keeping only those with the lowest 15% of all adjusted p-values and which are then significant in more than 50% of all the samples in each tissue over all ages.

### Pseudobulk and spatial neighborhood expression analysis

Expression analysis on the sample-level was performed using pseudobulk aggregation of normalized counts via the Seurat function AggregateExpression, either per experimental group or per spot cluster. Resulting count matrices were then scaled to zero mean and unit variance using the ScaleData function. Clustering of expression matrices was performed using hierarchical clustering with Euclidean distance metric. To perform a spot neighborhood expression analysis for parasite invasion hot spots in the infection cohort, we selected all spots with more than 0.5% of ANKA mapped reads, and pulled all up to six direct neighbors in the hexagonal spot array. We then compared the group of center spots against their neighbors using the Seurat *FindMarkers* function with a minimum log fold-change of 0.25, minimum percentage of expressed spots of 0.1, an alpha level of 0.05 as well as using Spot sample identity for latent variable regression. To analyze the combined expression of genes from the complement pathway via an activity score, we used the *AddModuleScore* function from Seurat with a fixed seed on the data slot of the spatial assay and all other parameters kept at default.

### Spatial neighborhood enrichment analysis

To perform spatial neighborhood analysis on our Visium 10x samples, we first converted all Seurat objects to Squidpy^101^ (v1.4.1) *AnnData* objects. This was accomplished by first extracting gene expression matrix, spatial coordinates and required metadata from the Seurat object, subsequently loading them into Squidpy using the *AnnData* function. To process groups of samples together, we first manually shifted the spatial coordinates of each sample such that there is no spatial overlap between samples in the same dimensions. Based on those common coordinates, we first calculated the spatial connectivity matrix using the function *gr.spatial_neighbours*. To validate that there is no inter-sample connectivity we created spatial scatter plots using the *pl.spatial_scatter* function with connectivity key set to *spatial_connectivities*. Then, we used the Squidpy function *gr.nhood_enrichment* with mode *zscore* to calculate the neighborhood enrichment matrix for each group of samples. To compare between two groups (e.g. young vs. old), we computed the pointwise difference between their neighborhood enrichment matrices, providing us with a differential neighborhood enrichment matrix per contrast.

### Pathway enrichment analysis

To perform gene set over-representation analysis on DEGs we used the Transcriptomics module of GeneTrail3^102^, with the target species set to Mus musculus, null hypothesis set to two-sided, and Benjamini-Hochberg FDR p-value correction at an alpha level of 0.05. We tested against Gene Ontology (Biological process, Cellular component, Molecular function), KEGG, Reactome, Wikipathway, and Pfam databases with all supported mouse genes as background normalization set.

### Analysis of small non-coding RNA sequencing data

Raw reads were cut, cleaned, filtered and mapped against small non-coding RNA annotations using the end-to-end pipeline miRMaster2^103^. Genome-wide mature miRNA counts were generated for each valid sample using the *Mus musculus* annotation of miRBase^104^ v22.1. Overall, three of 48 samples with unexpectedly low total read counts were removed from any further analysis. We then performed standard QC steps such as dimension reduction, sample and feature correlation and clustering analysis, count normalization as well as differential expression using our established pipeline in miRMaster2. Enrichment analysis was conducted using miEAA^105^ at default parameters and with all recommended category source databases enabled.

### Spatial metabolomics data generation

A timsTOF flex MALDI-2 (Bruker Daltonics) operated in positive MALDI (Matrix-Assisted Laser Desorption/Ionization) mode with Post Ionization laser and enabled ion mobility separation was used for the data acquisition. The Imaging 20 µm Smart Beam laser intensity was set to 85%, creating one burst of 200 Shots with a frequency of 1000 Hz per pixel. Trigger delay for the Post Ionization laser was set to 5 µs and the used laser power for the post ionization laser was calculated automatically. A scan range of 50-1300 *m/z* was chosen. The offset of the MALDI plate was set to 50 V, deflection 1 delta to 70 V, Funnel 1 RF to 250 Vpp, Funnel 2 RF and the multipole RF to 200 Vpp. The collision cell energy was set to 10 eV and the collision RF to 350 Vpp. The quadrupole low mass was set to 50 *m/z* with an ion energy of 5 eV. The transfer time for the pre TOF focus was set to 75 µs with a pre pulse storage of 5 µs. A mobility range given in 1/K0 was measured from 0.50 Vs cm^-2^ to 1.40 Vs cm^-2^ with a ramp time of 100 ms and an accumulation time of 193 ms. The duty cycle was set to 100% with a ramp rate of 5.02 Hz. Mass calibration was conducted with red phosphorus on the respective cluster formed with the MALDI laser and ion mobility calibration dimension was calibrated linearly using 4 selected ions from ESI Low Concentration Tuning Mix (Agilent Technologies, USA) [*m/z*, 1/k_0_: (322.048121, 0.7363 Vs cm^-2^), (622.028960, 0.9915 Vs cm^-2^), (922.0098, 0.9915 Vs cm^-2^)] in positive mode. The mobility values for the mobility calibration were taken from the CCS compendium.

Sample preparation including the slicing is identical to the transcriptomics sample preparation, using an embedding in 2% CMC (Carboxymethylcellulose) solution (water) and matched samples from the other brain hemispheres. The slices were transferred immediately onto ITO (Indium Tin Oxide) coated glass slides after cutting, dried at room temperature and put in a desiccator for 24 h. Before matrix application the edges of every tissue were marked with a solvent resistant pen by making four crosses in a rectangular order onto the ITO-coated slide in 2 mm distant from the tissue border each. Each glass slides was scanned to obtain a picture showing the position of the tissue and the marked crosses on the glass slide (EPSON perfection V30). 15 mg/mL 2,5 - Dihydroxybenzoic acid (DHB), dissolved in a mixture of acetonitrile and water (90:10) spiked with 0.1% TFA (trifluoroacetic acid) was sprayed onto the tissue (HTX M5 sprayer). Fourteen passes of matrix were added with a velocity of 1200 mm/min, a track spacing of 3 mm and a flow rate of 0.125 mL/min. The nozzle height was set to 40 mm and a temperature to 60°C while the nitrogen gas flow was set to 2 L/min and a pressure of 10 psi.

### Spatial metabolomics analysis

The automatic MALDI imaging runs were set up using fleximaging 7.6 (Bruker Daltonics) and the timsTOF flex MALDI-2 was operated with timscontrol 6.16. Raw data files were then processed together with mzmine^106^ v4.3.0 using batch mode and the following processing steps in order: 1) “Import MS data”, 2) “Mass detection”, 3) “Image builder”, 4) “Ims expander”, 5) “Smoothing”, 6) “Isotopic peaks finder”, 7) “Image co-localization”, 8) “Join aligner”, 9) “Feature list rows filter”, 10) “Local compound database search”, 11) “Export CSV”, 12) “Raw data export”. The resulting mass feature peak list was analyzed with MetaboAnalyst6.0^107^ and individual feature ion images were generated using the Bruker SCiLS lab software. MetaboAnalyst was configured to remove features with low variance using the interquartile range filter set to 40% and to remove features with low abundance using the mean intensity filter set to 40%. Peak intensities were then normalized using quantile normalization, log scaling and mean centering.

### Statistics and reproducibility

All scripts were executed in a reproducible environment using a custom Snakemake v7.32.4 workflow with dedicated conda environments, fixing each utilized R and python package to a particular version: R v4.3.2, python v3.12, numpy v1.26.4, pandas v2.2.1, tidyverse v.2.0.0, patchwork v1.2.0, umap v0.2.10.0, pals v1.8, colorspace v2.1_0, ggrepel v.0.9.5, complexupset v1.3.3, bioconductor-complexheatmap v.2.18.0, readxl v1.4.3, openxlsx v4.2.5.2, future v1.33.0, pbapply v1.7_2, bioconductor-mast v1.28.0, seurat v4.4.0, seuratobject v4.1.4, seurat-disk v0.0.0.9021, matrix v1.6_1.1, sctransform v0.4.1, bioconductor-glmgampoi v1.14.0, hdf5r v1.3.10, python leidenalg v0.10.2, leidenalg v1.1.3, reticulate v1.34.0, python-igraph v0.11.4, umap-learn v0.5.5, devtools v2.4.5, remotes v2.4.2, yaml v2.3.8, doparallel v1.0.17, data.table v 1.15.2, ggVennDiagram v1.5.2, viridis v0.6.5, ggdendro v0.2.0, fmsb v0.7.6, Stereopy (v1.1.0), SPARK-X (v1.1.1), and Squidpy (v1.4.1). Statistical tests were performed two-tailed if not stated otherwise. Raw p-values were adjusted for multiple testing bias to an alpha level of 0.05 using either the Bonferroni (DEG analysis) or Benjamini-Hochberg FDR (pathway analysis) correction method. Trend curves were obtained through Local Polynomial Regression Fitting (loess) of *y* ∼ *log*(*x*) with a span of 0.75, two degrees of freedom and a standard error confidence interval of 0.95.

## Data availability

The sequencing datasets and images generated throughout this study are available in the NCBI Sequence Read Archive (SRA) via accession-ID PRJNA1163328 (reviewer link: https://dataview.ncbi.nlm.nih.gov/object/PRJNA1163328?reviewer=43g3vvl2a3s5o0nnrksc23pu9) as well as Gene Expression Omnibus (GEO) via accession-ID GSE283283 (reviewer password: ybkzqyygdjatvad). Preprocessed Seurat R objects will also be made available upon request.

## Code availability

Custom scripts used to produce the results of this study are available from the corresponding authors upon request.

## Acknowledgements and funding

We thank our peers from all the contributing labs providing their continued support and feedback on the project. We are grateful for having received advanced user support and guidance from 10x Genomics and BGI on establishing their sequencing-based spatial transcriptomics platforms. This study was funded by the M.J. Fox Foundation (MJFF-021418), the Knight Initiative for Brain Resilience, the Schaller-Nikolich Foundation, the Hans-und-Ruth-Giessen Foundation, and Saarland University. Computational resources used within this study were financed through the DFG project 469073465. G.K-C. received further support by the DFG CRC TRR152 grant P22 and a DFG KR 4338/1-2 grant.

## Author contributions

F.K. and Vi.W. wrote the manuscript with input from all other authors; F.K. led the data analysis with support by S.G., J.A., A.H., M.F., F.G., and S.R.; Vi.W., Va.W, and N.L. performed the spatial transcriptomics experiments; A.F., M.A., B.K., A.H., H.L., T.J., and O.H. organized the cohort setup, animal treatment, organ extraction and sample logistics; Vi.W., Va.W, A.H., P.W., M.H., T.J., G.K-C., U.B., and O.H. supported the histological analysis and cluster annotation procedures; P.D., and D.K. performed the spatial metabolomics experiments and supported the data analysis; U.B., O.H., and T.W-C. supported the design of the mouse aging cohort; H.L., R.M., and T.J. supported the design of the mouse infection cohort; T.J., G.K-C., U.B., T.W-C., and A.K. supported data interpretation and supervised the study; F.K., T.W-C. and A.K. provided the study funding. All authors read and approved upon the manuscript. U.B., O.H., T.J., and G.K-C. contributed equally and were listed in alphabetical order of surnames.

## Supplementary Information

Supplementary Information is available for this paper.

**Supplementary Table 1**: Sample metadata and quality control metrics for the mouse aging cohort.

**Supplementary Table 2**: Spot cluster marker genes for peripheral organs from the aging cohort (10x Visium).

**Supplementary Table 3**: Aging DEGs per cluster and peripheral organ (10x Visium).

**Supplementary Table 4**: Aging SVGs per peripheral organ and sample (10x Visium).

**Supplementary Table 5**: Spot cluster marker genes for brain sections from the aging cohort (10x Visium).

**Supplementary Table 6**: Aging DEGs per cluster and brain bregma (10x Visium).

**Supplementary Table 7**: Aging SVGs per brain bregma and sample (10x Visium).

**Supplementary Table 8**: Spot cluster marker genes for brain sections from the aging cohort (STOmics Stereo-seq).

**Supplementary Table 9**: Aging DEGs per cluster and brain bregma (STOmics Stereo-seq).

**Supplementary Table 10**: Computationally aligned and annotated mass feature peaks for the aging brain spatial metabolomics experiments.

**Supplementary Table 11**: Sample metadata and quality control metrics for the mouse infection cohort.

**Supplementary Table 12**: Spot cluster marker genes for all organs from the infection cohort (10x Visium).

**Supplementary Table 13**: Infection DEGs per cluster and organ (10x Visium).

**Supplementary Table 14**: Infection SVGs per organ and sample (10x Visium).

**Supplementary Table 15**: Annotation, normalized counts, differential expression and enrichment statistics from miRNA sequencing of the infection cohort.

**Supplementary Table 16**: Raw GeneTrail3 results for the over-representation analysis of brain, kidney, liver and lung DEGs shared between aging and infection clusters.

## Extended Data Figures

**Extended Data Figure 1:**
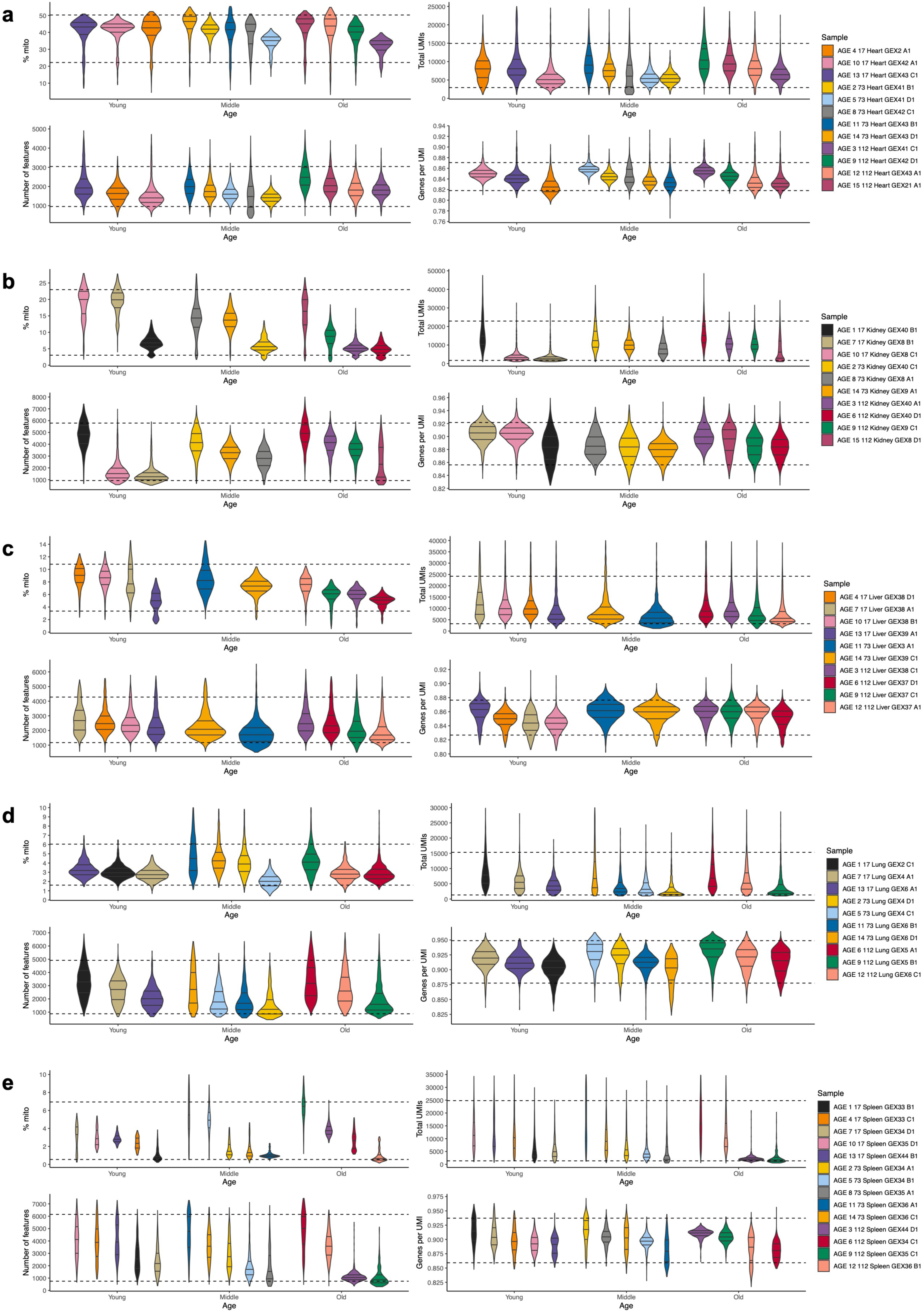
Aging peripheral organ samples quality control. **a-e**, Distribution of percentage of mitochondrial reads (top left), total number of UMIs (top right), number of detected gene features (bottom left) and spot complexity (bottom right), per cleaned high-quality sample and experimental group, for heart (**a**), kidney (**b**), liver (**c**), lung (**d**), and spleen (**e**).

**Extended Data Figure 2:**
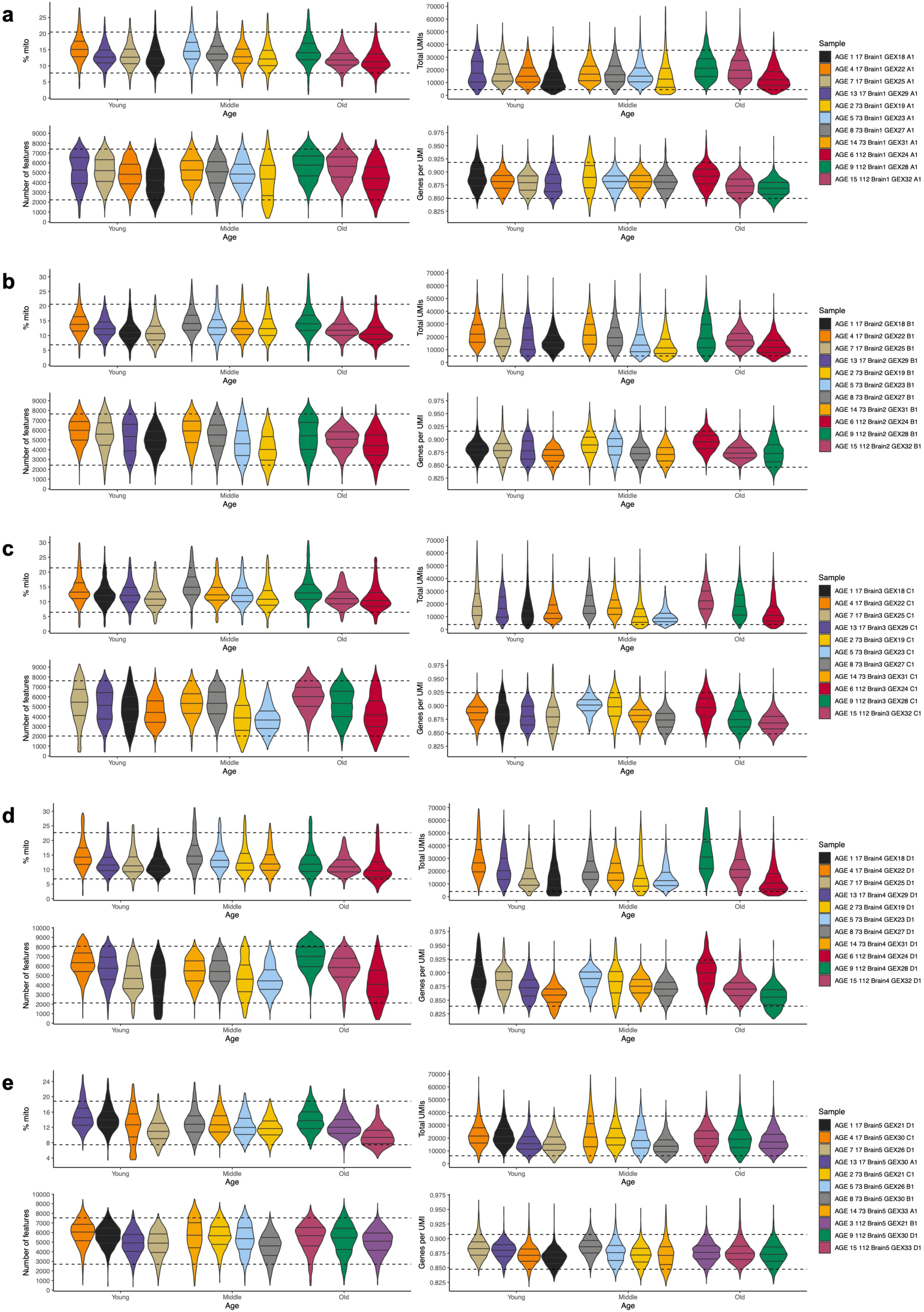
Aging brain samples quality control. **a-e**, Distribution of percentage of mitochondrial reads (top left), total number of UMIs (top right), number of detected gene features (bottom left) and spot complexity (bottom right), per cleaned high-quality sample and experimental group, for brain1 (**a**), brain2 (**b**), brain3 (**c**), brain4 (**d**), and brain5 (**e**).

**Extended Data Figure 3:**
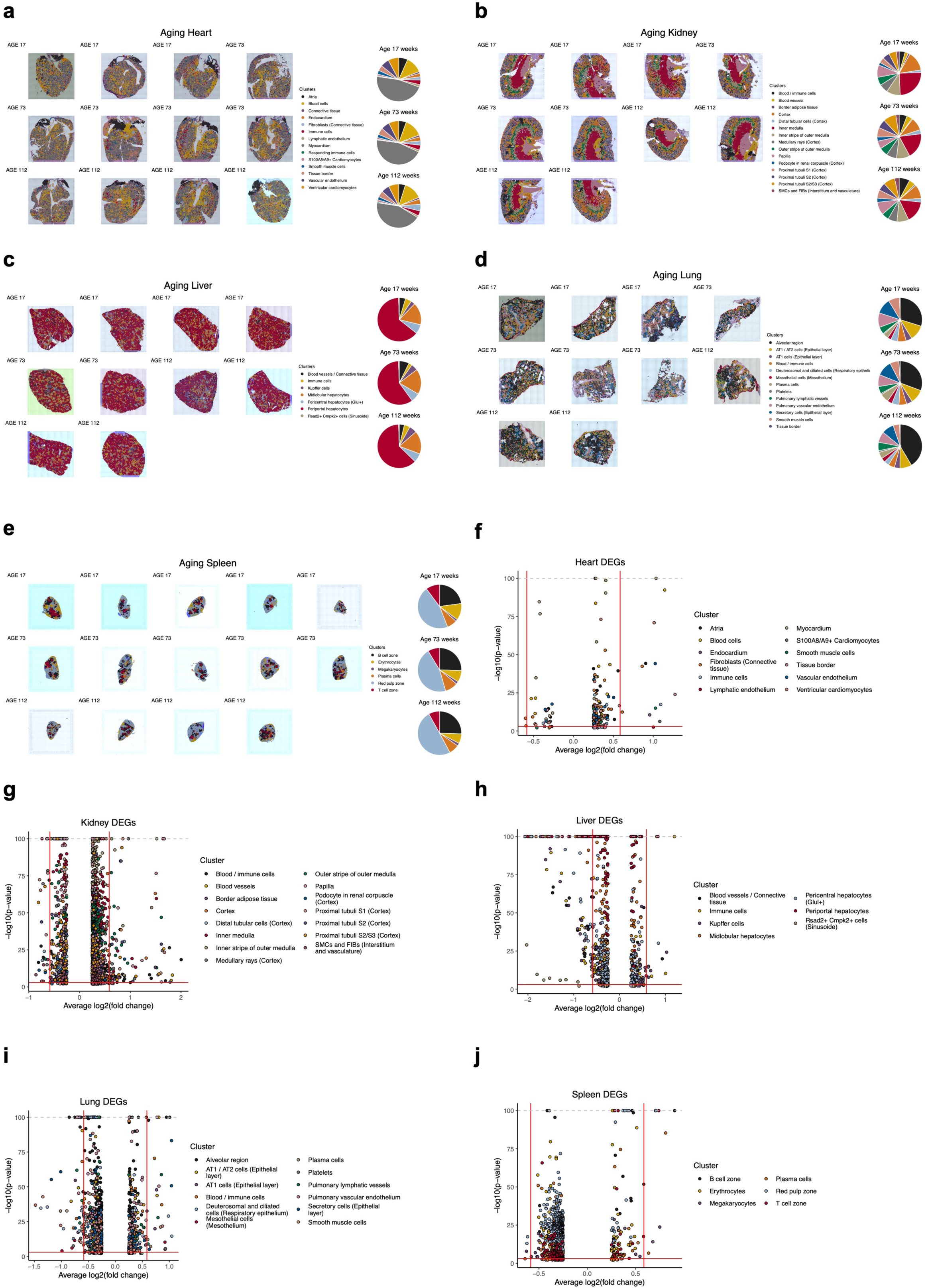
Integrated annotations and DEG analysis for the aging peripheral organs. **a-e**, H&E-stained image slices (left) ordered by age (top row: young, middle row: middle, bottom row: old) with annotated spot clusters overlaid for all high-quality samples from heart (**a**), kidney (**b**), liver (**c**), lung (**d**), and spleen (**e**). Pie charts (right) showing the proportion of spot clusters per age group. **f-j**, Volcano plots of aging DEGs (old vs. young) colored by spot cluster for heart (**f**), kidney (**g**), liver (**h**), lung (**i**), and spleen (**j**).

**Extended Data Figure 4:**
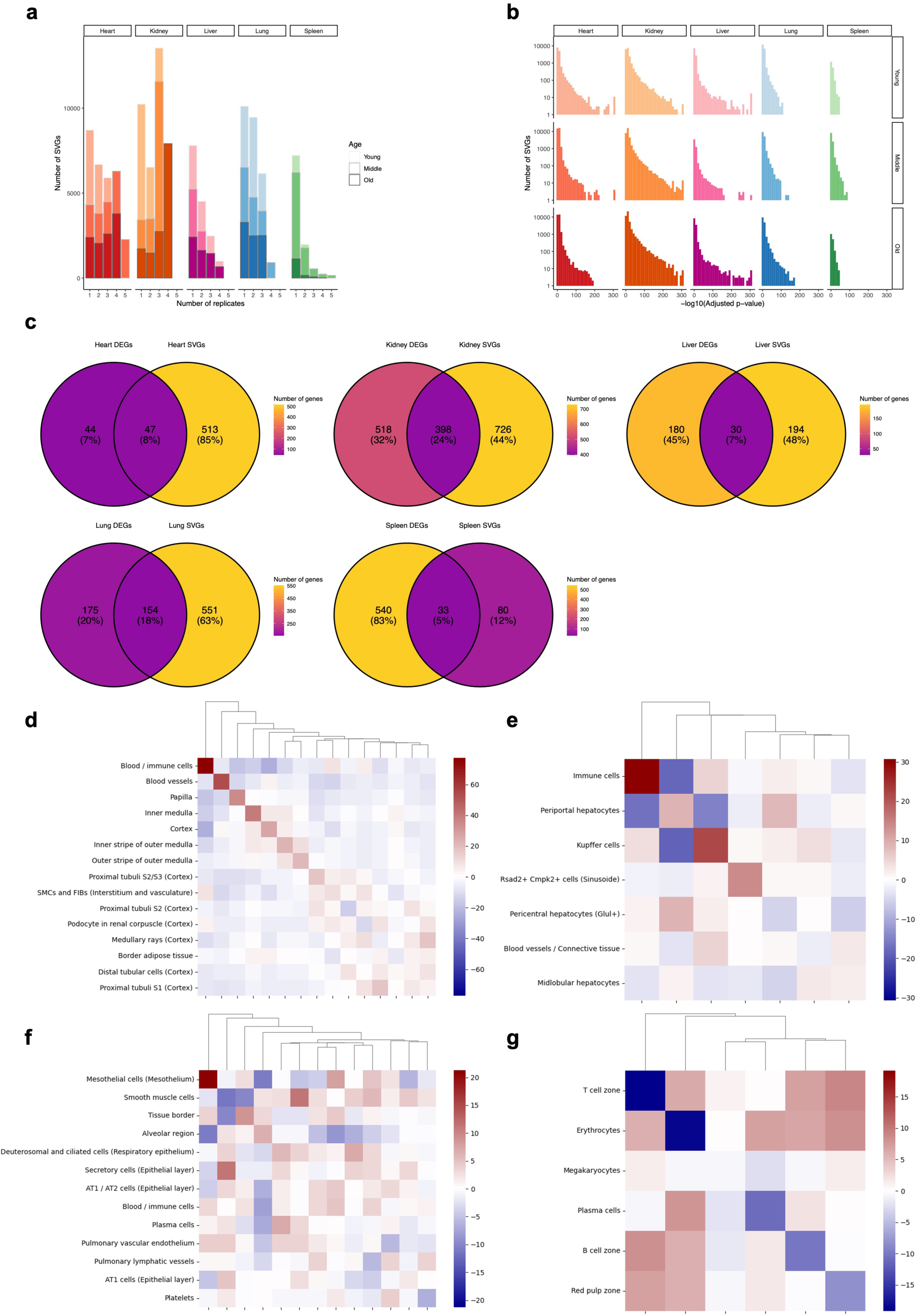
SVG analysis and neighborhood enrichments for aging peripheral organs. **a**, Bar plots showing the number of significant SVGs (y-axis) by the minimum number of replicate samples (x-axis) per age group and peripheral organ as determined with SPARK-X. **b**, Histogram showing the adjusted and -log10 scaled p-value distribution by age and peripheral organ for all significant spatially variables genes determined with SPARK-X. **c**, Venn diagram comparing the significant aging DEGs (old vs. young) and significant SVGs per peripheral organ. **d-g**, Like in Fig. 1j but for kidney (**d**), liver (**e**), lung (**f**), and spleen (**g**).

**Extended Data Figure 5:**
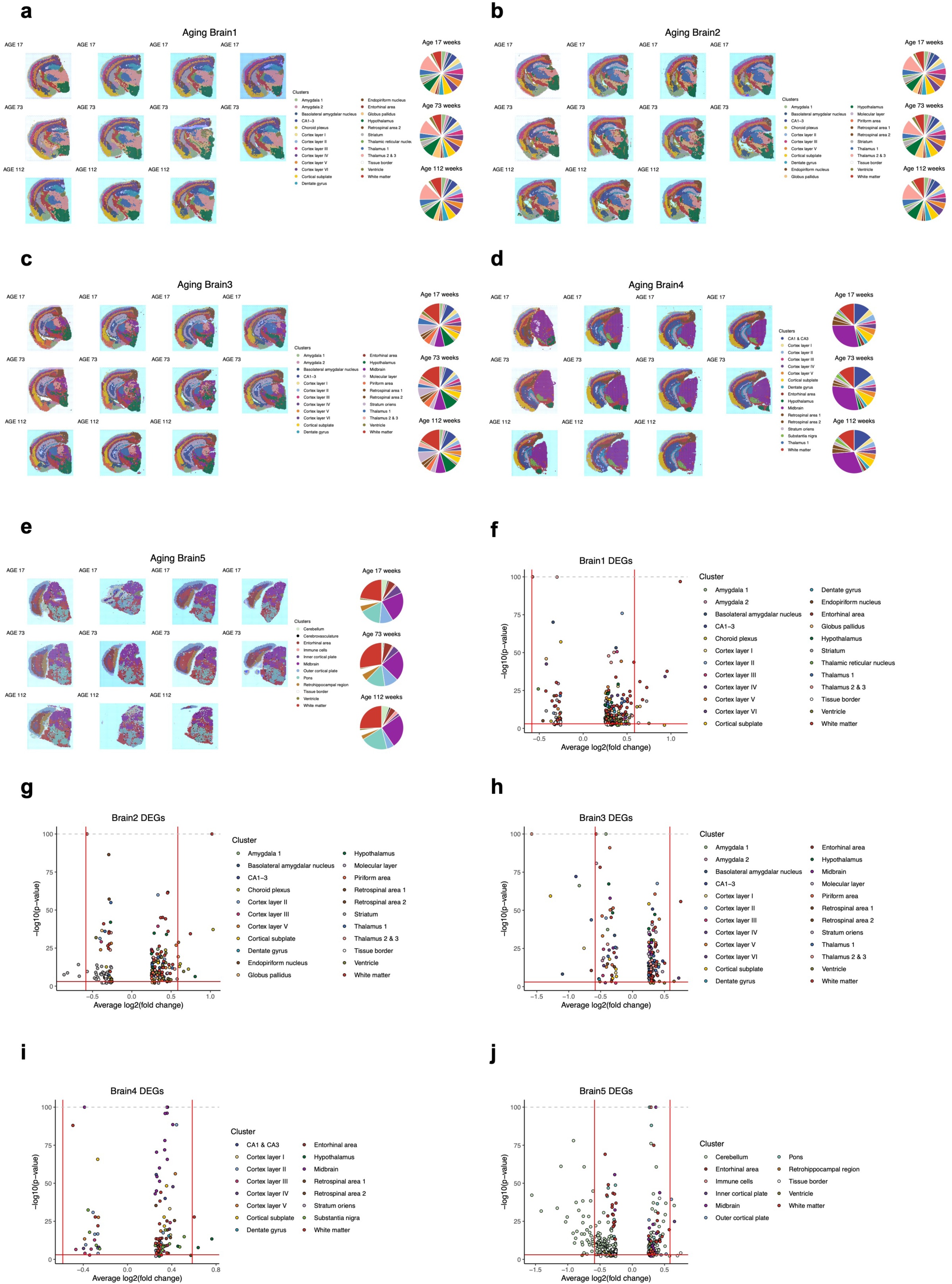
Integrated annotations and DEG analysis for the aging brain slices. **a-e**, As in **Extended Data Fig. 3a-e** but for the aging brain slices. **f-j**, As in **Extended Data Fig. 3f-j** but for the aging brain samples.

**Extended Data Figure 6:**
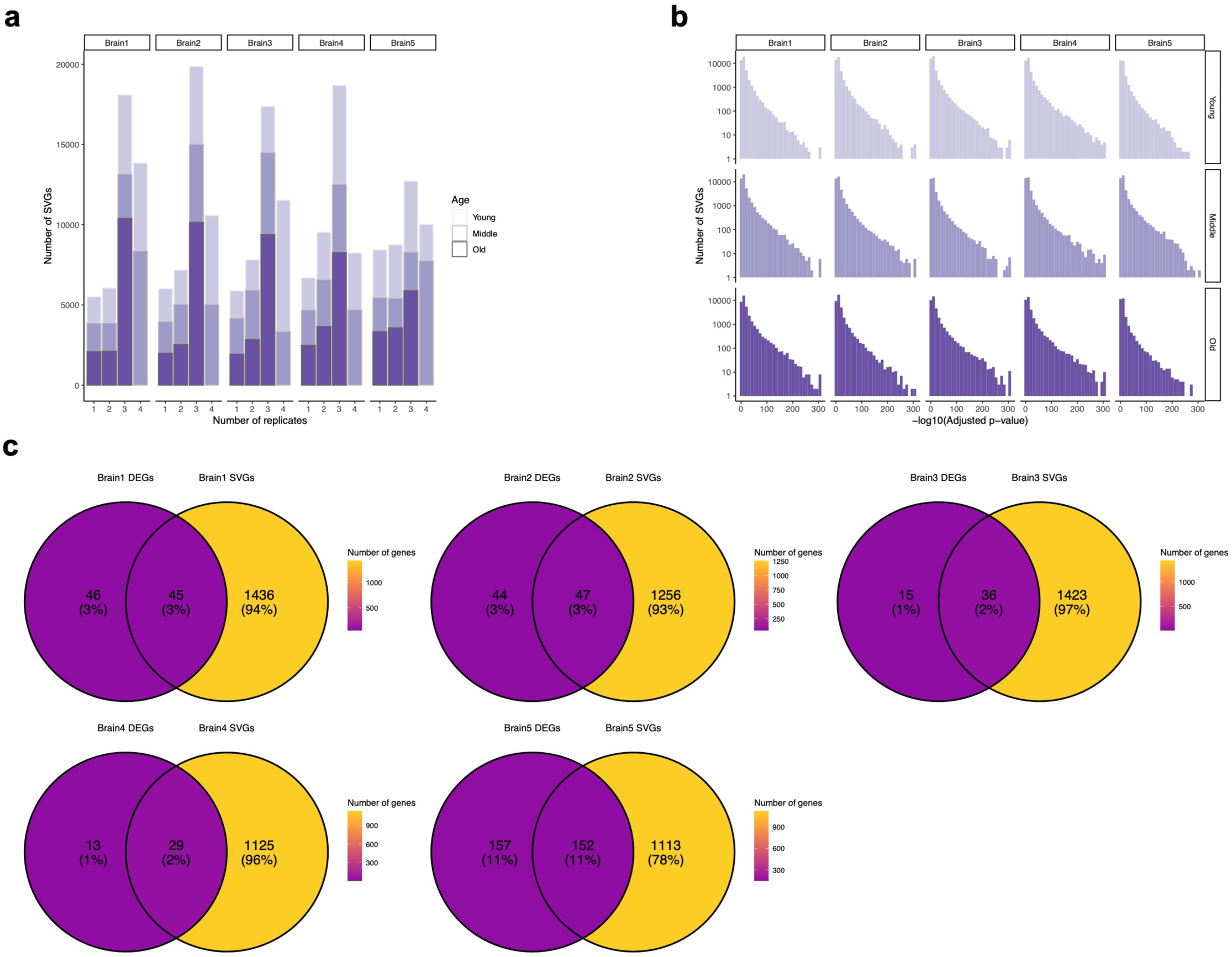
SVG analysis for aging brain slices. **a**, As in **Extended Data Fig. 4a** but for the aging brain samples. **b**, As in **Extended Data Fig. 4b** but for the aging brain samples. **c**, As in **Extended Data Fig. 4c** but for the aging brain samples.

**Extended Data Figure 7:**
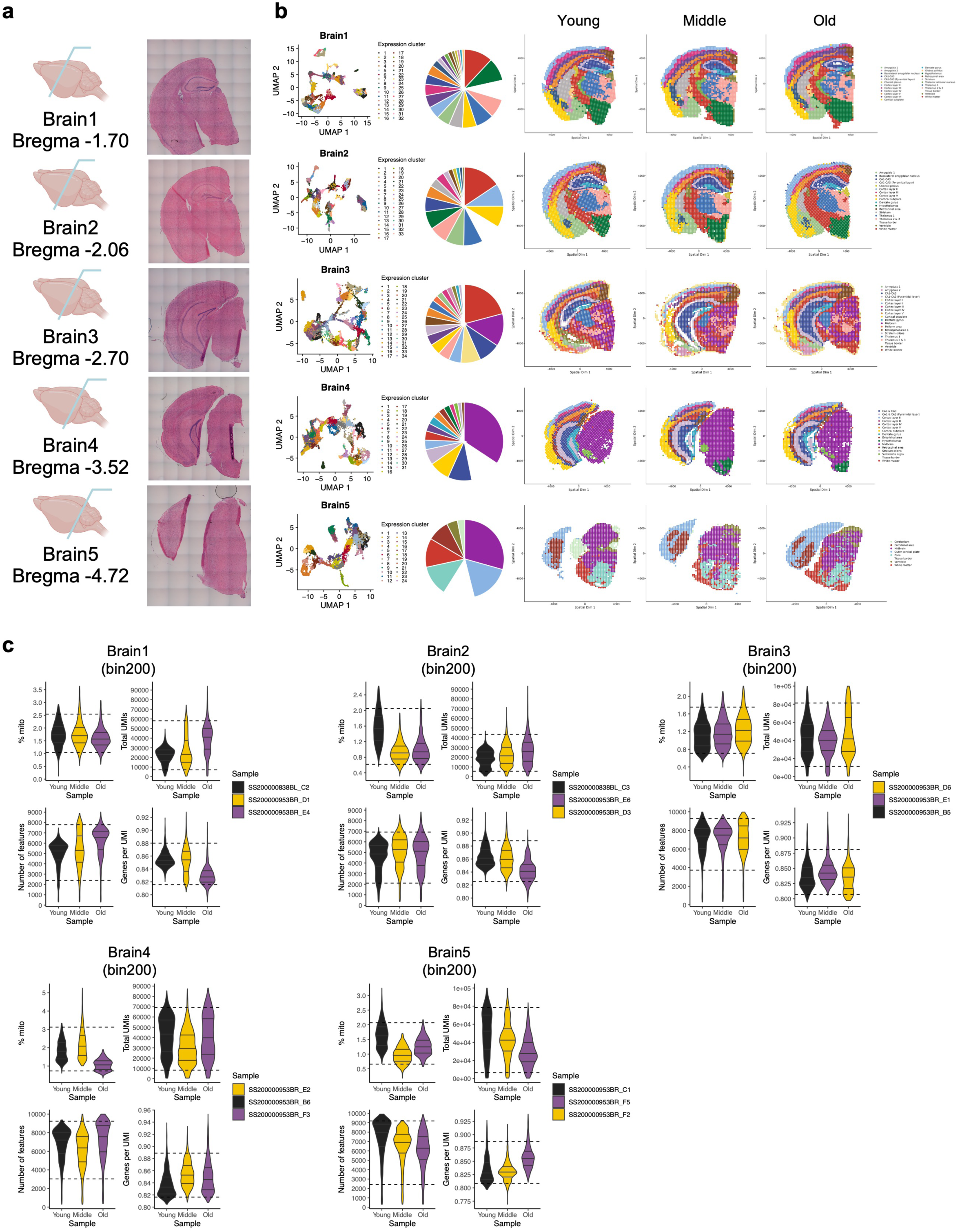
Quality control and annotations for STOmics aging brain slices. **a**, Illustration of the five different brain bregma used for STOmics Stereo-seq in accordance with the Visium data set. Representative H&E stains are shown for each Bregma. Since Stereo-seq does not support H&E stains directly from the sequenced tissue slices, an adjacent (directly before or after) tissue slice was prepared and stained before running the spatial transcriptomics experiments. **b**, From left to right and per brain bregma (top to bottom): integrated UMAP representation of all cleaned Stereo-seq spot clusters using the bin200 resolution, pie charts and per replicate spatial projections of the final annotated spot clusters. Cluster names and colors were assigned in accordance with the Visium data set (cf. Methods). **c**, Distribution of four main quality control features across the cleaned spots and per Stereo-seq brain replicate at bin200 resolution.

**Extended Data Figure 8:**
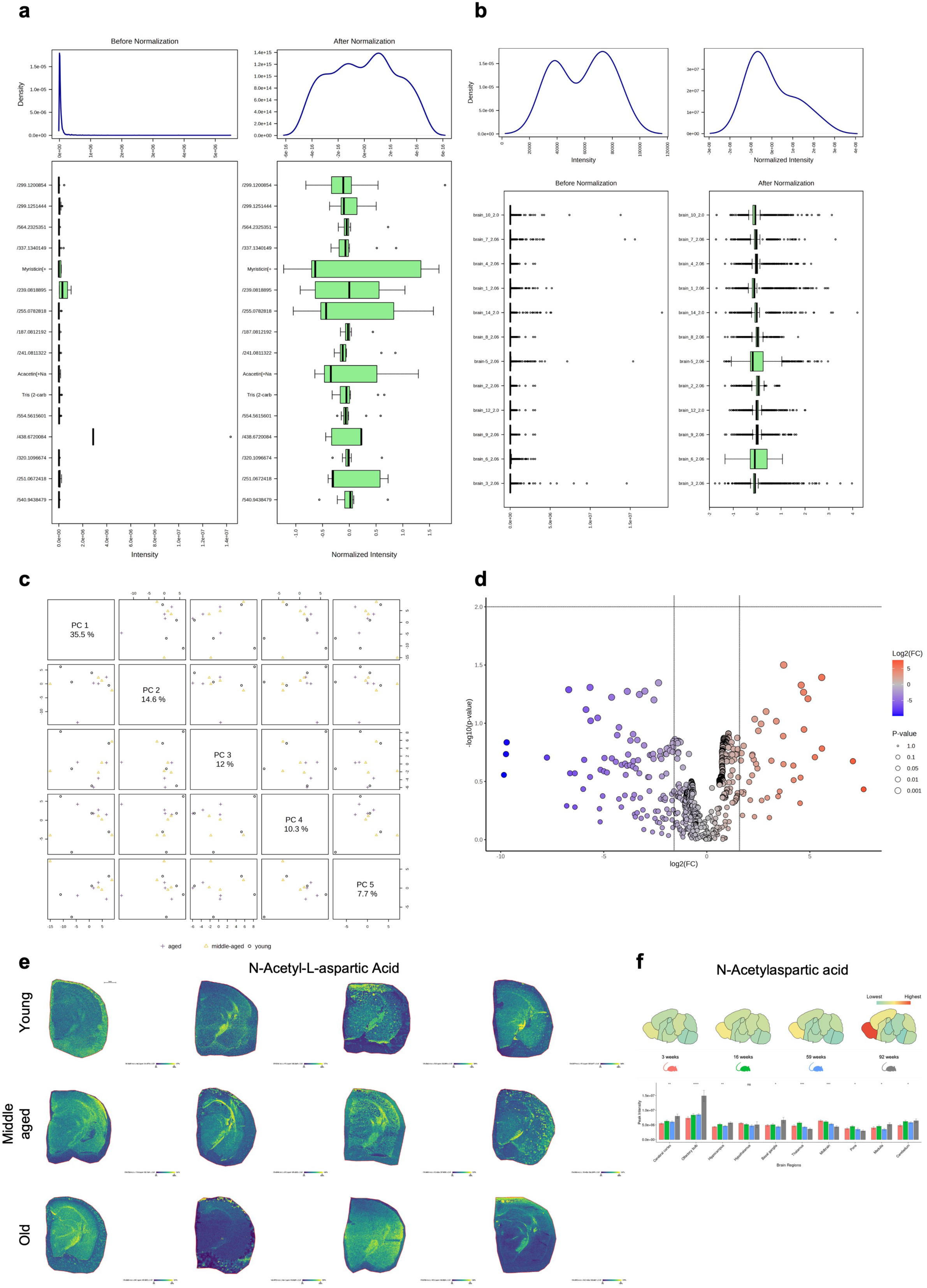
Spatial metabolomics of the aging brain. **a**, MS peak area intensities before and after normalization displayed by feature. **b**, MS peak area intensities before and after normalization displayed by sample. **c**, Two-dimensional principal component matrix for the first five dimensions from a PCA of peak areas after normalization. Point shape and color correspond to a sample age group. **d**, Volcano plot with raw unpaired t-test p-values against log fold-changes comparing spatial metabolomics peak features for old (N=4) versus young (N=4) mouse brains. Points are colored according to fold-change. **e**, Experimentally measured and computationally integrated peak intensities of N-Acetyl-L-aspartic Acid (∼ 176.0576 *m/z*) in the spatial domain of twelve brain hemispheres from the aging cohort. **e**, Peak intensities for N-Acetylaspartic acid across mouse brain regions and four age groups as reported by Ding et al^22^.

**Extended Data Figure 9:**
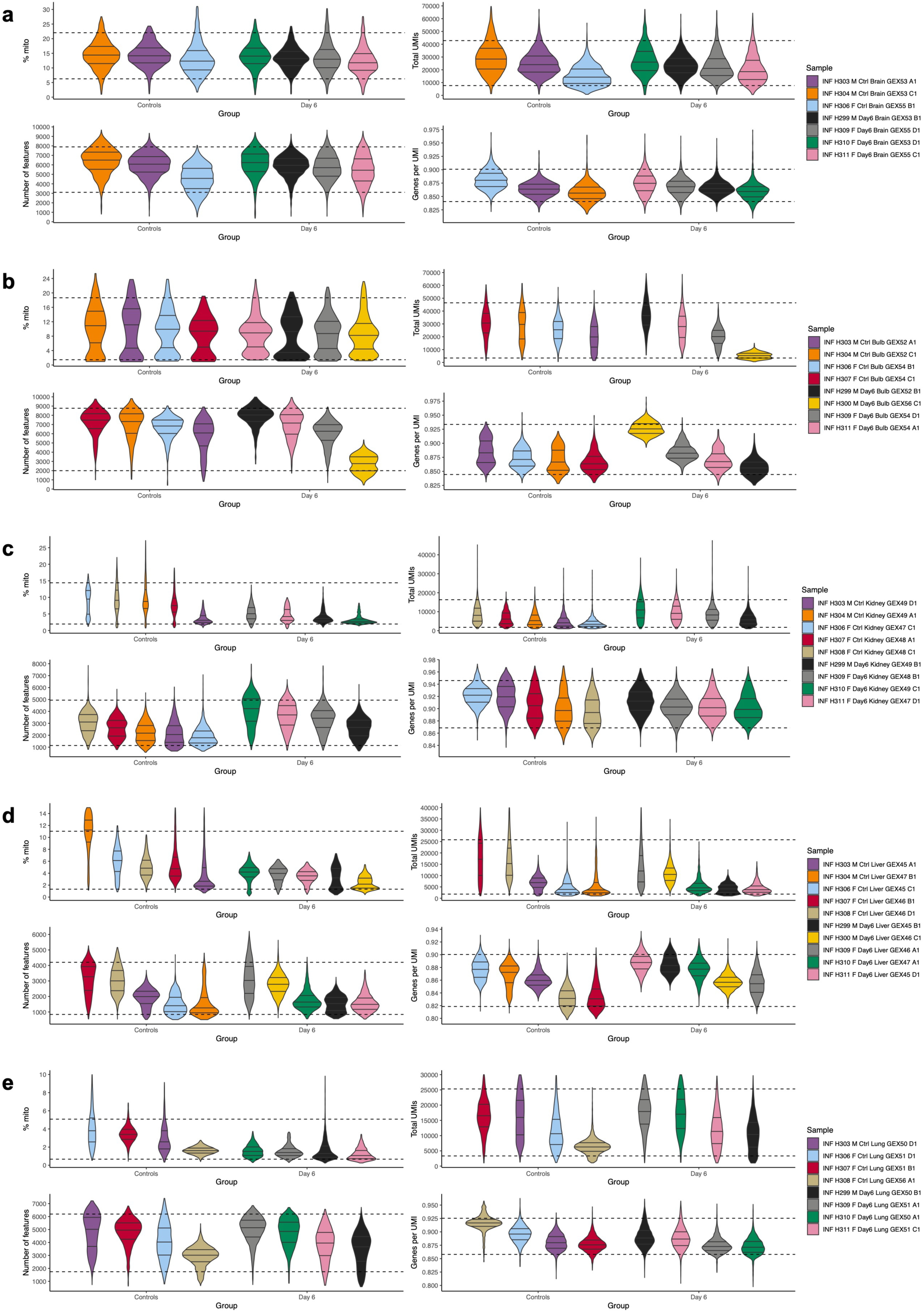
Quality control for organ samples of the infection cohort. **a-e**, Like in **Extended Data Fig. 1a-e** but for all organs from the infection cohort.

**Extended Data Figure 10:**
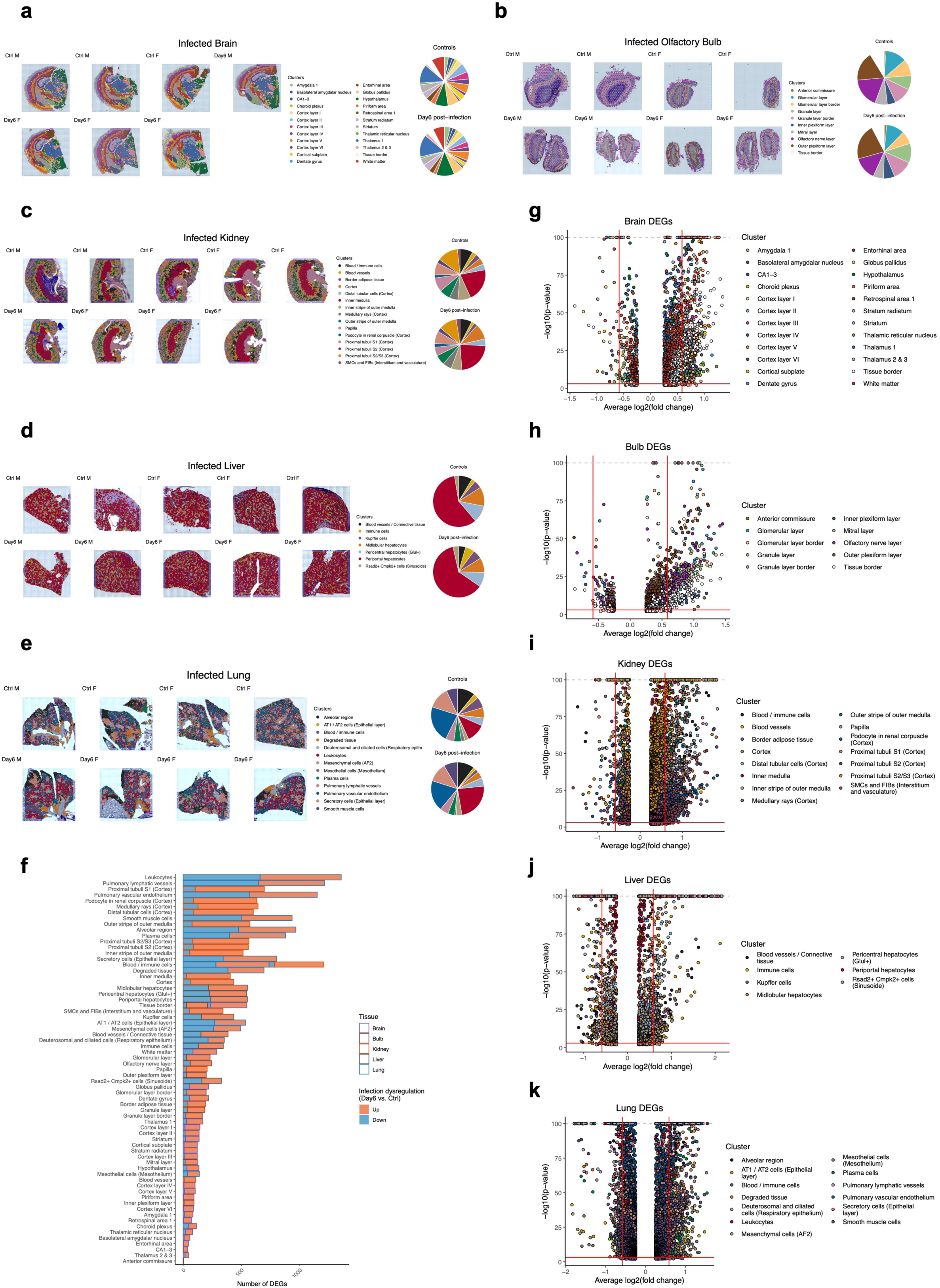
Integrated organ annotations and DEG analysis for the infection cohort. **a-e**, As in **Extended Data Fig. 3a-e** but for all the organs from the infection cohort, split by experimental group. **f**, Like in Fig. 1g, but for all tissue-cluster DEGs from the infection cohort. **g-k**, As in **Extended Data Fig. 3f-j** but for all the organs from the infection cohort, comparing day6 post-infection against healthy controls.

**Extended Data Figure 11:**
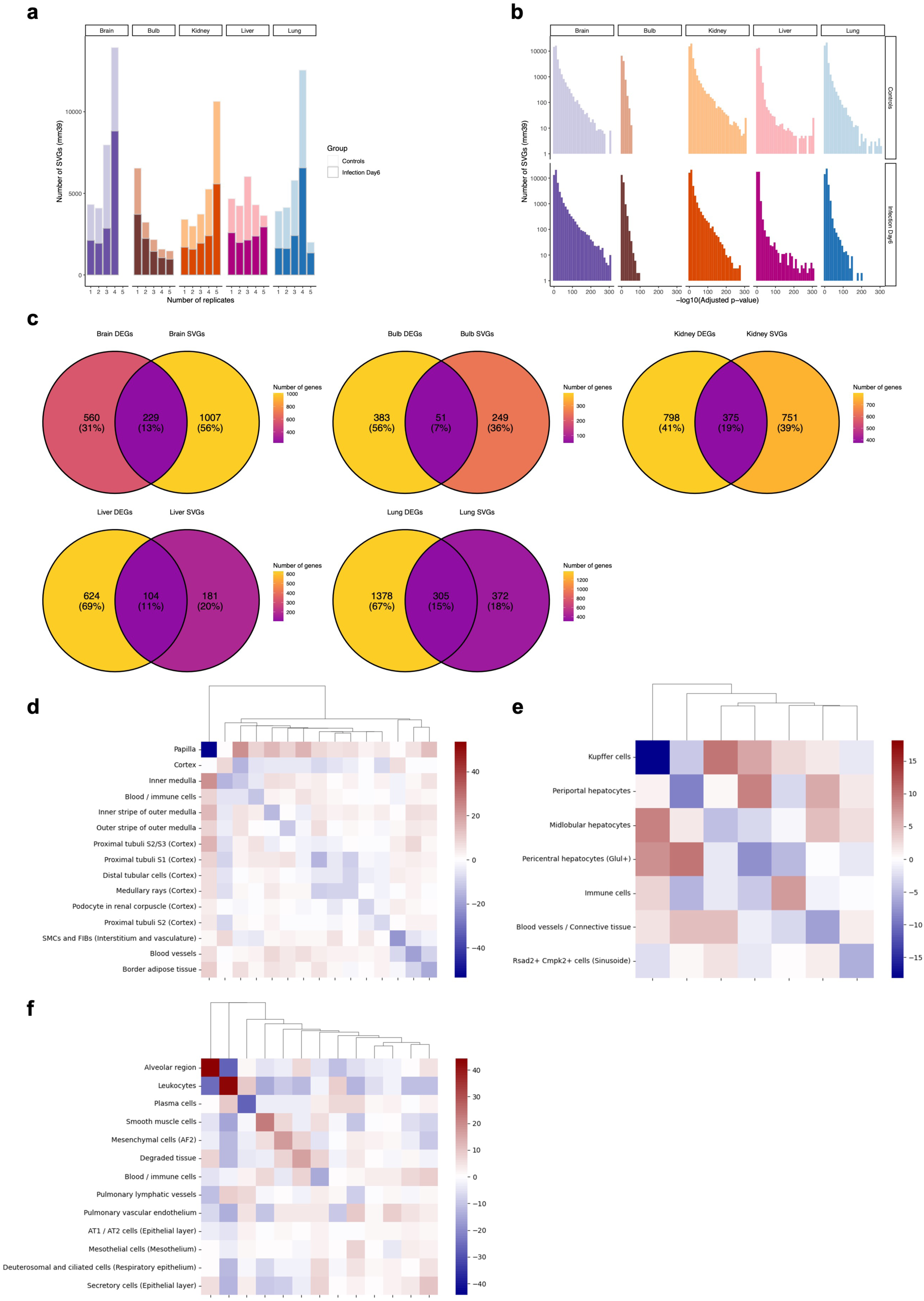
SVG analysis and neighborhood enrichments for the infection cohort. **a**, As in **Extended Data Fig. 4a** but for all the infection cohort organs, split by experimental group. **b**, As in **Extended Data Fig. 4b** but for all the infection cohort organs, split by experimental group. **c**, As in **Extended Data Fig. 4c** but for all the infection cohort organs, comparing SVGs against day6 post-infection vs. healthy control DEGs. **d-f**, Like in Fig. 1j but for kidney (**d**), liver (**e**), and lung (**f**) from the infection cohort.

**Extended Data Figure 12:**
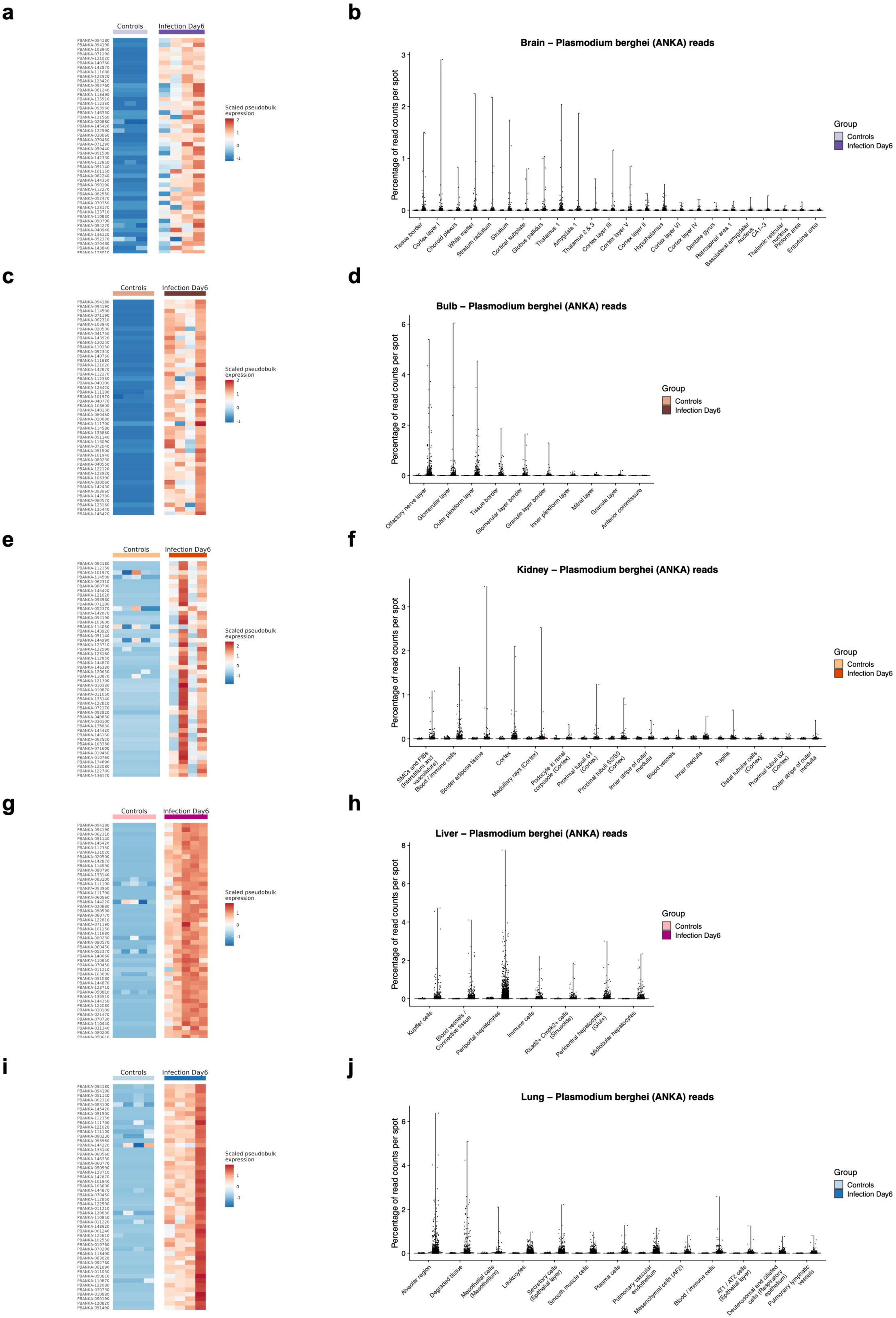
ANKA genome read detection analysis for the infection cohort. **a**, Heatmap showing the scaled pseudobulk expression by brain sample and experimental group of the top 50 most variable genes from the *P. berghei* (ANKA) genome. **b**, Distribution of the percentage of ANKA-related read counts per spot across all annotated spot clusters of the brain and split by experimental group. **c**, **d**, As in (**a**,**b**) but for olfactory bulb. **e**, **f**, As in (**a**,**b**) but for kidney. **g**, **h**, As in (**a**,**b**) but for liver. **i**, **j**, As in (**a**,**b**) but for lung.

**Extended Data Figure 13:**
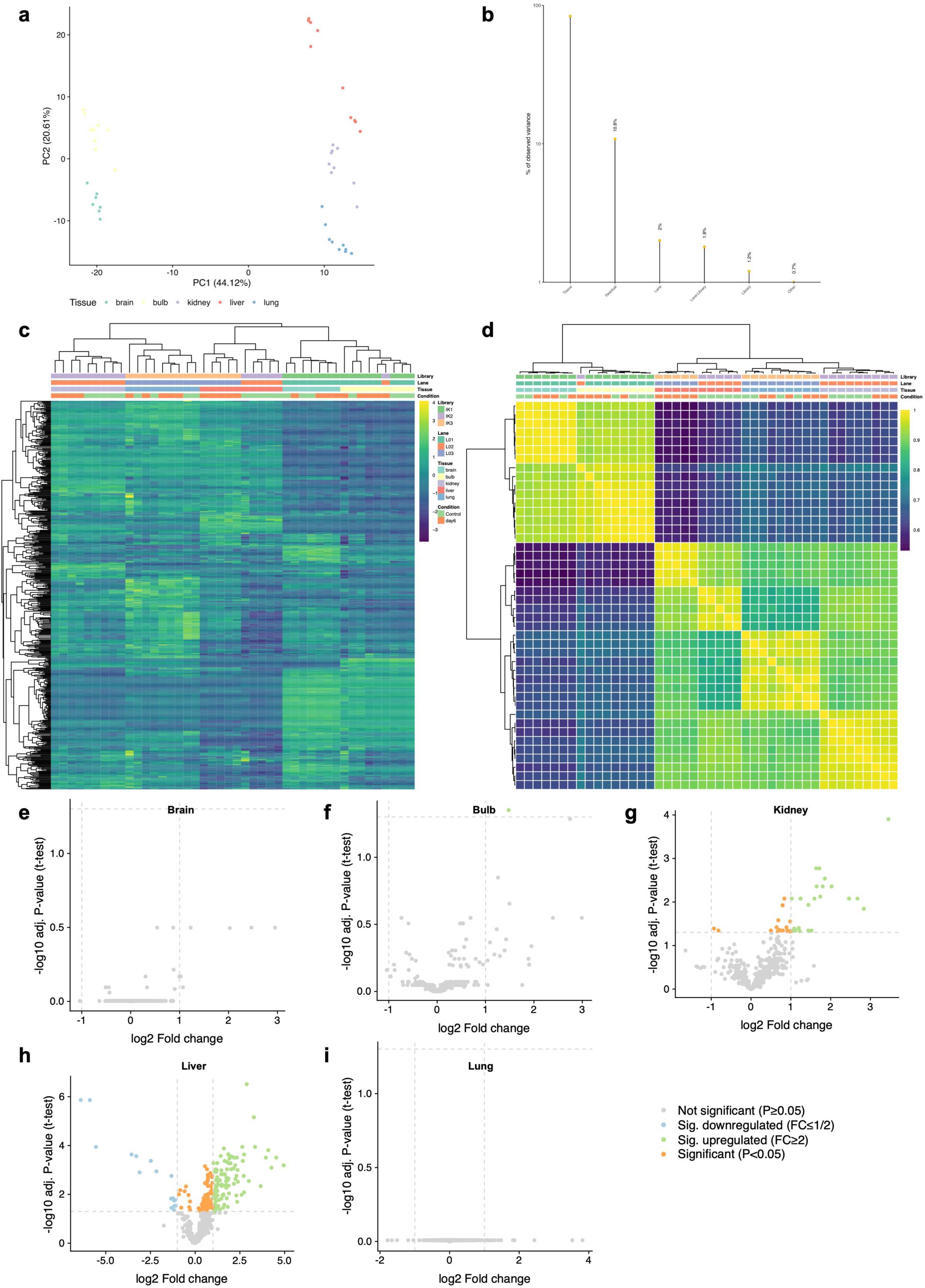
MicroRNA expression analysis for the infection cohort. **a**, Small non-coding RNA sequencing samples from the infection cohort projected onto the first two principal components of a PCA using the normalized miRNA expression. Each biological sample is a dot colored by tissue of origin. Values at axis labels describe the percentage of variance explained for each component. **b**, Results of the Principal Variance Component Analysis (PVCA) for the normalized miRNA expression from the infection cohort. Each bar shows the total percentage of explainable variance for a single covariate or a linear combination of such. The category ‘residual’ means the total variance left unexplained by all other variables. **c**, Heatmap of z-scores for the 542 miRNAs robustly detected in at least one organ for the infection cohort. The order of rows (miRNAs) and columns (samples) were determined using hierarchical clustering. **d**, Heatmap of sample-to-sample spearman correlation analysis. The order of rows (miRNAs) and columns (samples) were determined using hierarchical clustering. **e-i,** Volcano plots for miRNA differential expression statistics of brain, bulb, kidney, liver, and lung samples, comparing infected samples against controls.

**Extended Data Figure 14:**
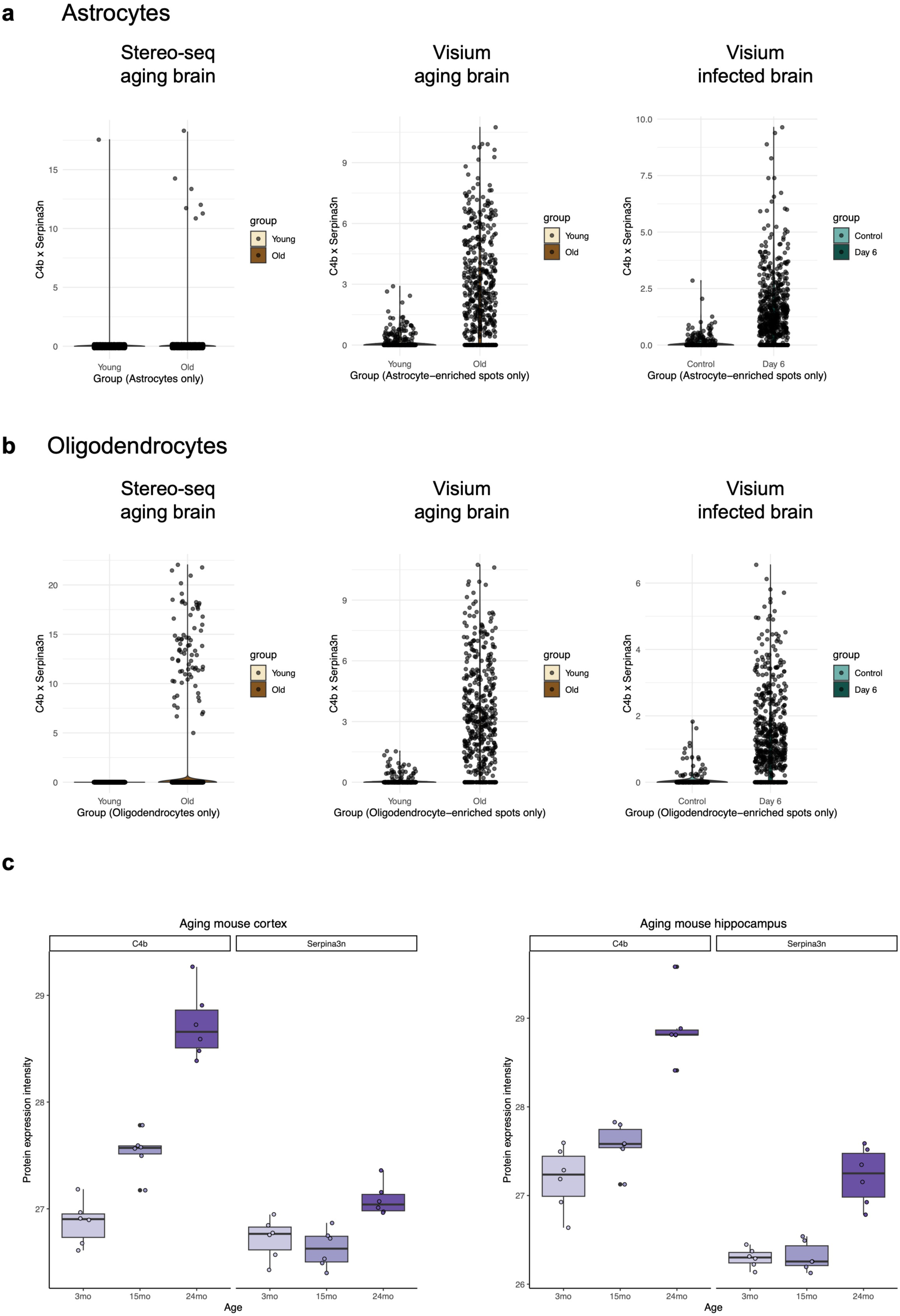
Validation of C4b and Serpina3n upregulation in the aging mouse brain. **a**, Combined analysis of C4b and Serpina3n by per-spot multiplication of normalized expression values across four groups of samples (young, old, control, infected). Shown are the combined expression values for Astrocyte assigned spots from the cell binning resolved brain1 Stereo-seq samples (left), and the Astrocyte marker enriched spots from the aging (middle) and malaria disease (right) mouse brain Visium samples. **b**, As in (**a**) but for the Oligodendrocyte assigned spots (Stereo-seq) and Oligodendrocyte marker enriched spots (Visium). **c**, Normalized protein expression intensities of C4b and Serpina3n in young (3 month), adult middle aged (15 month), and old (24 month) mouse brain cortex (left) and hippocampus (right), as originally obtained by Tsumagari *et al*^28^.

**Extended Data Figure 15:**
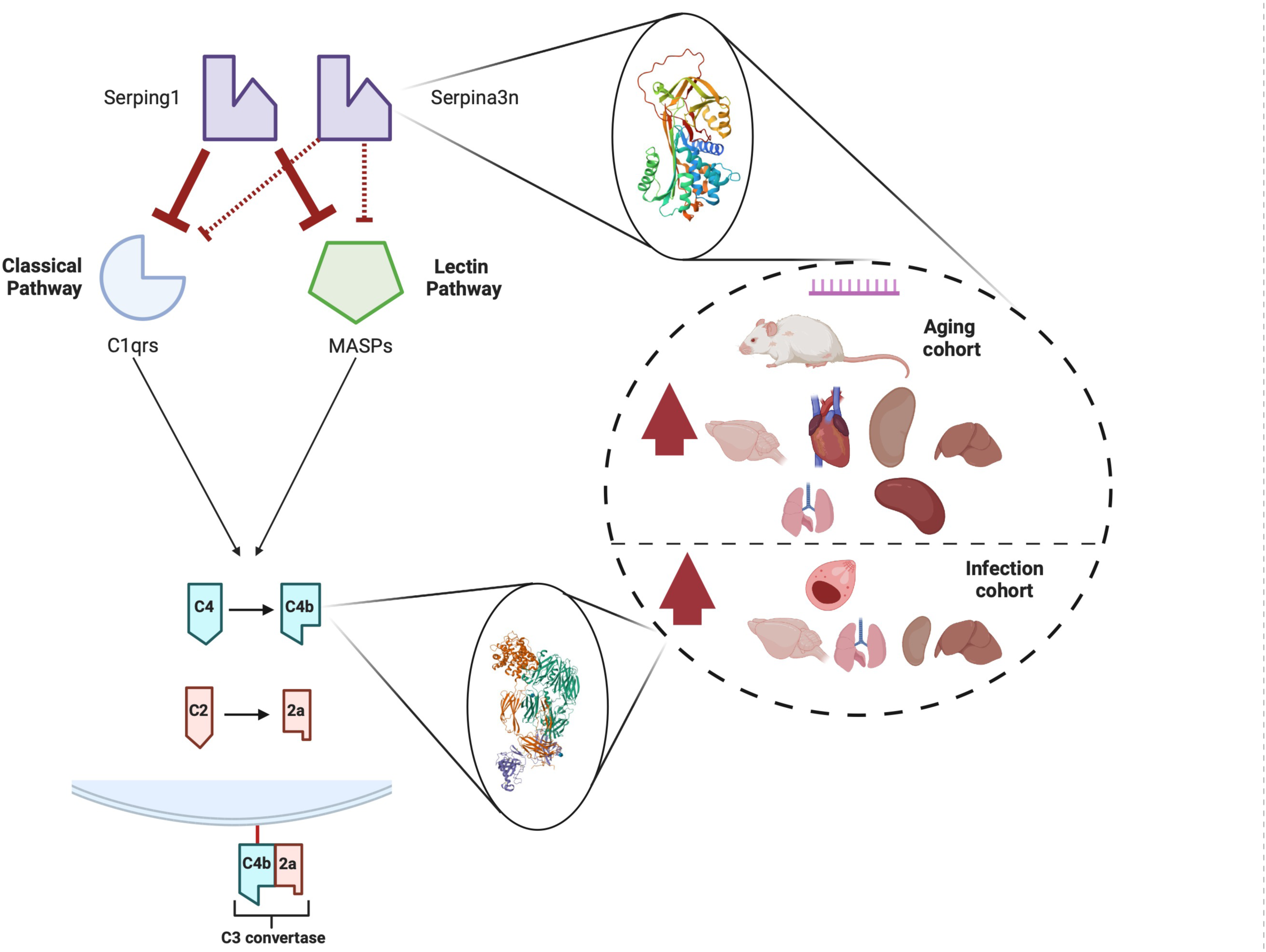
Summarized findings on complement and protease inhibitor activation. Members of the Serine protease inhibitor family regulate the expression of the classical and lectin complement pathways by interacting with individual proteases. We found a wide-spread and strong upregulation of these genes in the aging and malaria diseased mouse within different spatial compartments of several peripheral organs and the central nervous system (e.g. white matter tract), a gene signature which can also be traced down to individual cell types.

